# αKG inhibits Regulatory T cell differentiation by coupling lipidome remodelling to mitochondrial metabolism

**DOI:** 10.1101/2020.08.24.256560

**Authors:** Maria I. Matias, Carmen S. Yong, Amir Foroushani, Erdinc Sezgin, Kandice R. Levental, Ali Talebi, Cédric Mongellaz, Jonas Dehair, Madeline Wong, Sandrina Kinet, Valérie S. Zimmermann, Ilya Levental, Laurent Yvan-Charvet, Johannes V. Swinnen, Stefan A. Muljo, Saverio Tardito, Valérie Dardalhon, Naomi Taylor

## Abstract

The differentiation of CD4 T cells to a specific effector fate is metabolically regulated, integrating glycolysis and mitochondrial oxidative phosphorylation (OXPHOS) with transcriptional and epigenetic changes. OXPHOS is tightly coordinated with the tricarboxylic acid (TCA) cycle but the precise role of TCA intermediates in CD4 T cell differentiation remain unclear. Here we demonstrate that α-ketoglutarate (αKG) inhibited regulatory T cell (Treg) generation while conversely, increasing Th1 polarization. In accord with these data, αKG promoted the effector profile of Treg-polarized chimeric antigen receptor-engineered T cells against the ErbB2 tumor antigen. Mechanistically, αKG significantly altered transcripts of genes involved in lipid-related processes, inducing a robust lipidome-wide remodelling and decreased membrane fluidity. A massive increase in storage and mitochondria lipids was associated with expression of mitochondrial genes and a significantly augmented OXPHOS. Notably, inhibition of succinate dehydrogenase activity, the bridge between the TCA cycle and the electron transport chain, enforced Treg generation. Thus, our study identifies novel connections between αKG, lipidome remodelling and OXPHOS in CD4 T cell fate decisions.

## Introduction

The potential of a T cell to respond to infections, tumor antigens, and even auto-antigens is dependent on a massive augmentation of cellular resources, with the energetic profile of the cell intricately related to the harnessing of these resources (reviewed in (Almeida et al., 2016; Geltink et al., 2018; Makowski et al., 2020; Yong et al., 2017a)). The energetic state of the cell is also linked to the extracellular environment as activation of the T cell receptor (TCR) leads to an increase in the transcription and translation of nutrient transporters, resulting in an augmented uptake of the corresponding nutrients (Cretenet et al., 2016; Hukelmann et al., 2016; Macintyre et al., 2014; Sinclair et al., 2013). Indeed, incorporation of glucose- and glutamine-derived carbons into nucleotides promotes optimal T cell proliferation (Buck et al., 2017; Carr et al., 2010; Clerc et al., 2019; Klysz et al., 2015; Loftus and Finlay, 2016; Wang et al., 2011).

Nutrient resources do not merely provide energy and substrates for nucleotide synthesis; their availability and manner of utilization regulates the specificity of the T cell response to a foreign stimulus. This regulation is especially critical in the context of CD4 T lymphocytes where effector cells are highly glycolytic, and even lipogenic, while suppressive regulatory T cells (Tregs) display a mixed metabolism with increased levels of lipid oxidation (reviewed in (Buck et al., 2017)). Indeed, genetic deletion of nutrient transporters that are responsible for the uptake of glucose, leucine or glutamine, inhibits the differentiation of CD4 T cells to an effector fate (Teff) but not to a regulatory T cell fate (Macintyre et al., 2014; Nakaya et al., 2014; Sinclair et al., 2013). Moreover, limiting nutrient resources negatively impacts Teff function (Cham et al., 2008; Chang et al., 2013; Klysz et al., 2015; Macintyre et al., 2014; Metzler et al., 2016; Nakaya et al., 2014) while conversely, increasing glycolysis results in enhanced effector function, monitored as a capacity to produce IFNg (Buck et al., 2017; Chang et al., 2013; Peng et al., 2016). This balance in Teff/Treg differentiation is critical in the context of a tumor environment where the competition between T cells and tumor cells for limiting amounts of nutrients has a negative impact on the former.

The energetic profile of an activated CD4 T cell is dependent on an intricate crosstalk between various nutrient sensing pathways. mTOR, integrating extracellular and intracellular growth signals, has been identified as a critical regulator of CD4 T cell differentiation; mTOR inhibition blocks the generation of Th effector cells, promoting the generation and function of FoxP3-expressing Tregs (Delgoffe et al., 2009; Patel and Powell, 2017; Sun et al., 2018). Furthermore, alterations in metabolic sensors such as HIF-1α (Bottcher et al., 2018; Dang et al., 2011; Shi et al., 2011), inhibition of *de novo* fatty acid synthesis (Berod et al., 2014), and induction of a glycolytic metabolism over mitochondrial oxidative phosphorylation (OXPHOS) have all been shown to abrogate Th17 differentiation while promoting a Treg fate (Gerriets et al., 2015; Shin et al., 2020).

The balance between Tregs and Th1 effector cells secreting high levels of IFNg is also regulated by the extracellular metabolic environment; as compared to Th1 cells, Tregs are less impacted by reduced glycolytic capacity (Beier et al., 2015; Gerriets et al., 2015), functioning in a low-glucose, lactate-rich environment (Angelin et al., 2017). Furthermore, Th1 differentiation, but not Treg differentiation is dependent on extracellular glutamine (Klysz et al., 2015; Metzler et al., 2016). Interestingly though, mitochondrial metabolism plays a role in the function of both Treg and Th1 cells— mitochondrial integrity as well as complex III activity of the electron transport chain (ETC) are required for Treg suppressive activity (Beier et al., 2015; Field et al., 2019; Miska et al., 2019; Weinberg et al., 2019) whereas complex II activity is a requirement for terminal Th1 function (Bailis et al., 2019). Consistent with a role for OXPHOS in Th1 activity, the catalytic conversion of glutamine to alpha-ketoglutarate (αKG), which directly fuels the tricarboxylic acid cycle (TCA), has been shown to play a critical role in Th1 cell differentiation (Johnson et al., 2018), with αKG directly rescuing Th1 differentiation under glutamine-deprived conditions (Klysz et al., 2015). Importantly though, the role of αKG is complex, also functioning as an epigenetic modulator; αKG alters DNA and histone methylation states (Chisolm et al., 2017) as well as the specific methylation of the FoxP3 gene locus (Xu et al., 2017). Indeed, TET demethylases, which are dependent on αKG for the conversion of 5-methylcytosine (5mC) to 5-hydroxymethylcytosine (5hmC) (Lio and Rao, 2019; Rose et al., 2011; Tahiliani et al., 2009), are required for Treg stability and immune homeostasis (Nakatsukasa et al., 2019; Yang et al., 2015; Yue et al., 2019; Yue et al., 2016). However, inhibiting the generation of αKG, and thereby attenuating the reduction of αKG to the 2-hydroxyglurate (2-HG) competitive TET inhibitor (Xu et al., 2011), has also been shown to skew naïve T cells from a Th17 towards a Treg fate (Xu et al., 2017). Thus, the overall contributions of αKG to Treg differentiation are not yet completely understood.

Here, we show that under conditions of increased αKG levels, Treg polarization of naïve CD4 T was significantly diminished and inflammatory cytokines were induced. Conversely, the polarization of naïve CD4 T cells to a Th1 fate was significantly augmented, with a greater than 15-fold increase in levels of IFNγ transcripts. Consistent with these data, αKG promoted the effector profile of chimeric antigen receptor (CAR)-engineered T cells against the ErbB2 tumor antigen in tumor-bearing mice, increasing tumor infiltration and IFNγ secretion under conditions of Treg polarization. Notably, gene expression profiling of CD4 lymphocytes activated under Treg polarizing conditions in the presence of high levels of αKG revealed a significant enrichment of genes mechanistically associated with lipid processes. These transcriptional changes were coupled to a lipidome-wide remodelling, particularly associated with a significantly increased incorporation of polyunsaturated fatty acids (PUFAs) into membrane lipids. Furthermore, a >2-fold increase in storage lipids and mitochondrial lipids, together with a marked enrichment of mitochondrial respiratory genes, was coupled to a significantly increased energetic profile. We found that high intracellular levels of TCA cycle intermediates and oxidative phosphorylation directly regulated Treg differentiation. Inhibition of succinate dehydrogenase (SDH), also known as mitochondrial complex II—the bridge enzyme between the TCA cycle and the ETC (Rasheed and Tarjan, 2018), by either malonate or 3-nitropropionic acid (3-NP) (Huang et al., 2006; Quastel and Wooldridge, 1928) significantly augmented Treg polarization. Our data reveal a critical role for non-oxidative lipid metabolism in T cell fate--αKG induces a lipidome-wide remodelling and downstream OXPHOS that act in concert to establish equipoise between Th1 and Treg differentiation.

## Results

### αKG inhibits Treg polarization while conversely enhancing Th1 differentiation

As the potential of a CD4 T cell to generate a suppressive Treg phenotype is often inversely related to its potential to adopt an effector Th1 phenotype (Delgoffe et al., 2011; Klysz et al., 2015; Zeng et al., 2013), we assessed whether αKG levels modulate the balance between Treg and Th1 differentiation. While the majority of naïve T cells stimulated under Treg polarization conditions upregulated the master transcription factor FoxP3, the percentage of FoxP3^+^ T cells differentiated in the presence of exogenous αKG was significantly lower, decreasing from a mean of 60±2% to 28±2% (p<0.0001; **Figure 1A**). This decrease in Treg polarization was associated with 3.4-fold lower FoxP3 mRNA levels (p<0.05) and an attenuation of RNASeq reads in the *Foxp3* gene (**Figure 1B**). Moreover, Treg differentiation was reduced even when T cells were initially pre-stimulated in control Treg polarization conditions and αKG was only added 24h post-activation (Control, 68±8%; αKG, 24±4%; αKG at 24h, 35±9%; p<0.05; **Supplemental Figure 1**). Importantly though, the αKG-mediated inhibition of FoxP3 upregulation was specific to Treg polarizing conditions because the activation of thymic-Treg (tTreg), expressing FoxP3 in a quiescent state, was not altered by αKG (78±4% and 75±4%, respectively, **Figure 1C**).

**Figure 1:**
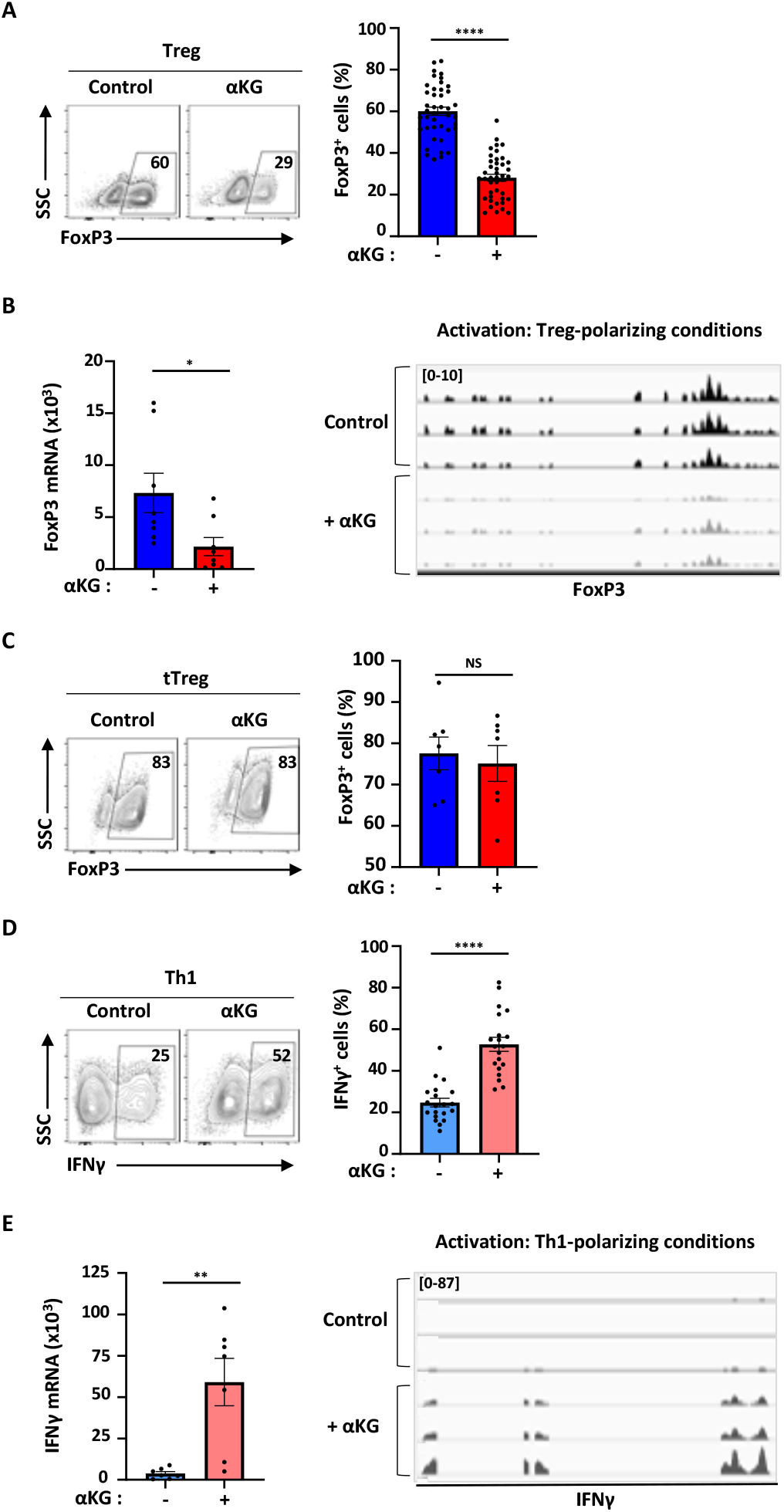
αKG augments Th1 polarization and inhibits the polarization of naïve CD4 T cells towards a Treg fate. (**A**) Naïve CD4 T cells were stimulated with α-CD3/α-CD28 mAbs under Treg-polarizing conditions in the absence or presence of αKG and the percentage of FoxP3^+^ Tregs was evaluated at day 4. Representative dot plots are shown (left) and quantifications of 42 independent experiments are presented (right). Means ± SEM are shown. (**B**) FoxP3 mRNA levels were evaluated by qRT-PCR and normalized to HPRT. Each point represents an independent experiment (left, n=8) and means ± SEM are shown. RNAseq read densities of the *Foxp3* gene in Treg-polarized samples in the absence or presence of αKG are presented (n=2 independent experiments performed with technical triplicates, right panels). (**C**) FoxP3^+^ thymic Tregs (tTregs) were isolated from FoxP3-GFP reporter mice and activated with α-CD3/α-CD28 mAbs in the absence or presence of αKG. At day 4, the percentages of FoxP3^+^ cells were evaluated by flow cytometry (left) and quantification of means ± SEM of 7 independent experiments are presented (right). (**D**) Naïve CD4 T cells were stimulated under Th1-polarizing conditions in the absence (control) or presence of exogenous αKG. The percentages of IFNγ^+^ cells were evaluated by intracellular staining and representative dot plots showing the percentages of IFNγ^+^ cells are presented (left). Quantification of 20 independent experiments are shown with means ± SEM presented (right panels). (**E**) IFNγ mRNA levels were assessed by qRT-PCR at day 4 of Th1 polarization and normalized to HPRT. Each point represents an independent experiment (left panel, n=7). Genome browser shots of RNA-seq reads over the *Ifng* gene in Th1-polarizing cells in the absence or presence of αKG are shown with the range of reads per million (RPM) presented on the y axis (n=3, right panels). All statistical analyses were evaluated by an unpaired 2-tailed t-test (*, p<0.05; **, p<0.01; ****, p<0.0001; NS, non-significant).

Conversely, αKG significantly augmented the percentage of CD4 T cells producing IFNγ, increasing the mean level from 25±2 to 53±3% (p<0.0001; **Figure 1D**). This increase was regulated at the transcriptional level as we detected an approximate 16-fold augmentation in IFNγ mRNA levels (p<0.01) and a marked increase in RNASeq reads mapped to the *Ifng* gene (**Figure 1E**). Thus, high levels of αKG augment the potential of CD4 T cells to produce IFNγ following stimulation in Th1-polarizing conditions. Together, these data reveal the importance of αKG in regulating the balance in the polarization of naïve CD4 T cells to a Th1 versus a Treg fate.

### αKG reprograms Treg towards an inflammatory phenotype

As αKG decreased the potential of CD4 T cells to differentiate into Treg, it was of interest to evaluate the consequence of reduced FoxP3 upregulation by this metabolite. Interestingly, the potential of these TGF-β-induced CD4 cells to produce IFNγ was significantly augmented by αKG, increasing from 0.5±0.1% to 5.8±0.6% (p<0.0001; **Figure 2A**) and this correlated with an upregulation of the master Th1 transcription factor, T-bet, at both the protein and mRNA level (**Figure 2B**). Furthermore, these CD4 T cells harbored higher levels of granzyme B (GzmB) transcripts (**Figure 2C**), a marker of CD4 T cells with cytotoxic T lymphocyte (CTL) activity (Takeuchi and Saito, 2017). It is also notable that the increased intracellular IFNγ levels (**Figures 2A**) paralleled the rise in IFNγ transcripts (**Figure 2D**). While neither TNF nor GM-CSF transcripts were increased, secretion of IFNγ, IL-17A, as well as TNF and GM-CSF were markedly elevated following stimulation of naïve CD4 T cells in Treg-polarizing conditions in the presence of αKG (p<0.01 to p<0.0001; **Figure 2E**). Together, these data highlight the inflammatory nature of these CD4 T cells, despite their activation under Treg-polarizing conditions.

**Figure 2:**
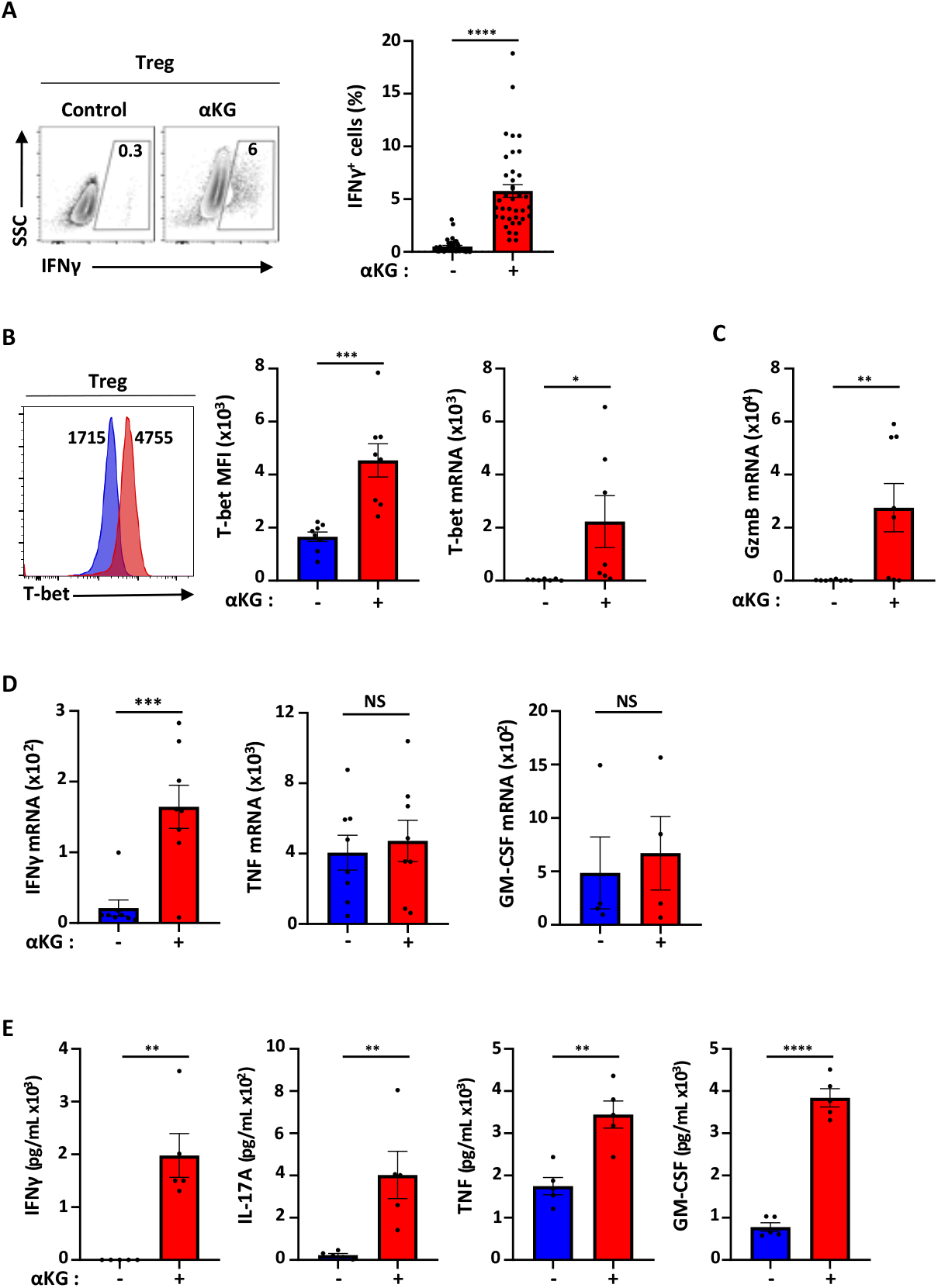
αKG induces an inflammatory profile in CD4 T cells activated under Treg-polarizing conditions. **(A)** Naïve CD4 T cells were activated under Treg polarizing conditions in the absence or presence of αKG and the percentages of IFNγ^+^ T cells were assessed at day 4 by intracellular staining. Representative dot plots (left) as well as quantifications of means ± SEM of 40 independent experiments (right) are shown. **(B)** T-bet protein levels were evaluated by flow cytometry and representative histograms are shown (left) as well as a quantification of the mean fluorescent intensity (MFI) ± SEM of 8 individual experiments (right). T-bet transcripts were quantified by qRT-PCR and normalized to HPRT (n=7) **(C)** Granzyme B (GzmB) transcripts were quantified by qRT-PCR and normalized to HPRT. Each point shows an independent experiment with means ± SEM presented (n=7). **(D)** mRNA levels for IFNγ (n=8), TNF (n=8) and GM-CSF (n=4) were assessed by qRT-PCR and normalized to HPRT. Each point represents data from an independent experiment. **(E)** Cytokine secretion was quantified at day 3 of Treg polarization by cytometric bead array (before the addition of supplementary IL-2). IFNγ, IL-17A, TNF, and GM-CSF levels in 5 independent experiments are presented with means ± SEM shown. All statistical analyses were evaluated by an unpaired 2-tailed t-test (*, p<0.05; **, p<0.01; ***, p<0.001; ****, p<0.0001; NS, non-significant).

### αKG-modulation enhances tumor infiltration of ErbB2-CAR T cells in mice bearing ErbB2^+^ tumors

The data presented above suggested that αKG might alter the phenotype and persistence of anti-tumor T cells in an *in vivo* setting. This is particularly critical since it has been shown that the intratumoral presence of Tregs negatively impacts the potential of effector T cells to eradicate tumor (Darrasse-Jeze et al., 2009) and the “hostile” environment of the tumor consumes high levels of nutrients (Yong et al., 2017b). To test this hypothesis, we evaluated whether αKG may affect the effector function of transgenic murine T cells harboring a chimeric antigen receptor (CAR) directed against the ErbB2/HER2 tumor antigen (ErbB2-CAR) (**Figure 3A**). Efficacy was evaluated against the 24JK fibrosarcoma, engineered to express ErbB2 (24JK-ERB), in RAG2^-/-^ mice (Yong et al., 2016). First, we found that similarly to polyclonal T cells, αKG treatment of transgenic ErbB2-CAR CD4 T cells resulted in a ≈2-fold increase in IFNγ secretion following activation in Th1 conditions and a decrease in FoxP3^+^ cells upon activation in Treg conditions (**Supplemental Figure 2**).

**Figure 3:**
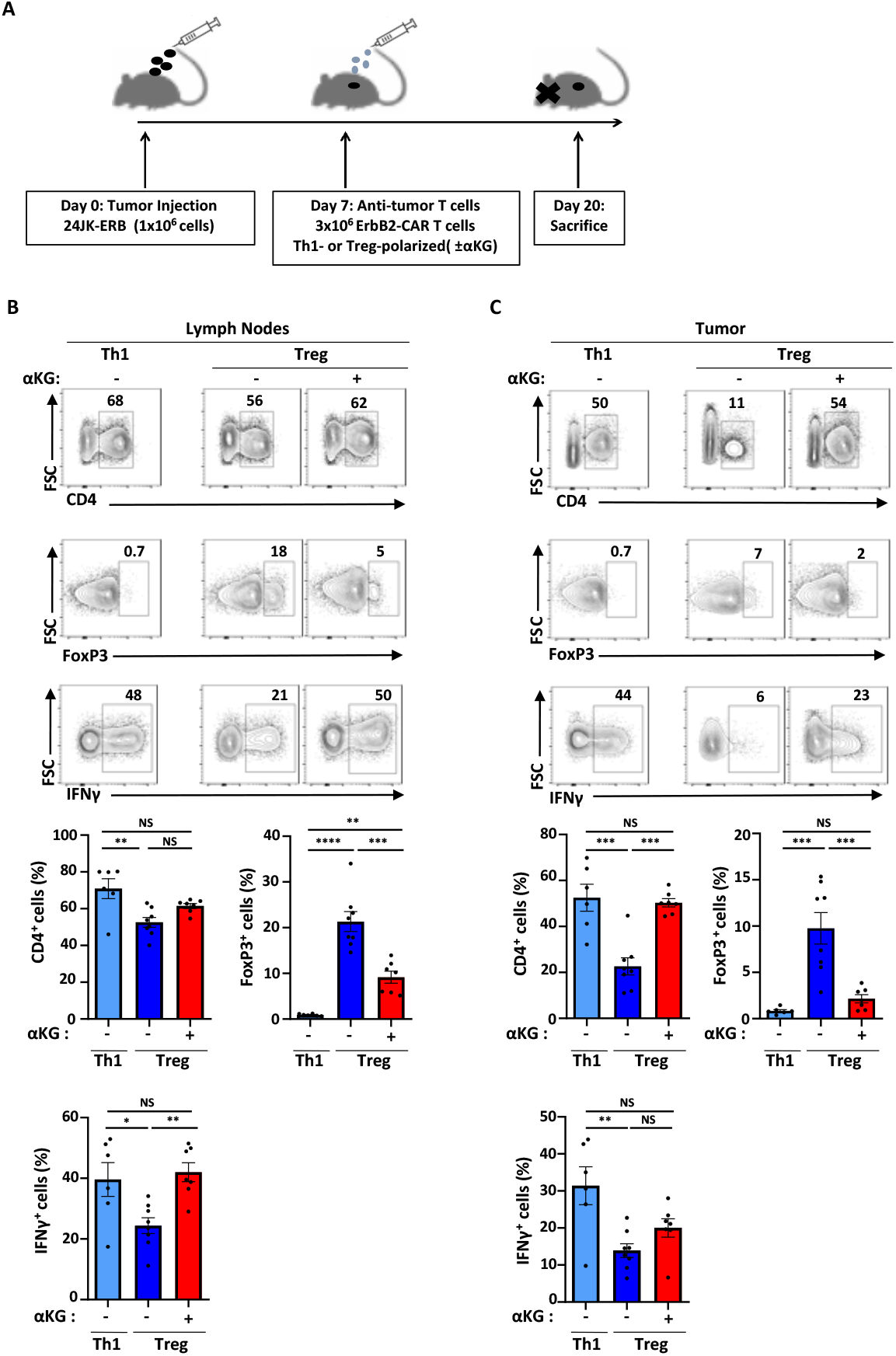
ErbB2-CAR T cells polarized *ex vivo* in the presence of αKG maintain an *in vivo* inflammatory profile following adoptive transfer into tumor-bearing mice. **(A)** Schematic of the experimental setup evaluating the impact of ErbB2-CAR T cells in mice bearing ErbB2^+^ tumors. RAG2^-/-^ mice were subcutaneously injected with ErbB2^+^ 24JK fibrosarcoma (24JK-ERB) and at day 7, CD4^+^ ErbB2-CAR T cells activated in either Th1 or Treg polarizing conditions, in the absence or presence of αKG, were adoptively transferred. Mice were sacrificed 13 days later (day 20). **(B)** The percentages of CD4^+^ as well as FoxP3^+^ and IFNγ^+^ CD4^+^ T cells recovered from draining lymph nodes (dLN) of tumor-bearing mice and representative plots are shown (top). Quantifications of the percentages of CD4^+^ T cells with the indicated phenotypes are presented with each point representing data from an individual mouse from 2 independent experiments (bottom, n=6-8). **(C)** The percentages of CD4^+^ as well as FoxP3^+^ and IFNγ^+^ CD4^+^ T cells recovered from tumors were evaluated. Representative plots (top) and quantifications (bottom) are presented as above. Statistical differences were determined by a one-way ANOVA and Tukey test for multiple comparisons (*, p<0.05; **, p<0.01; ***, p<0.001; ****, p<0.0001; NS, not-significant).

Naive ErbB2-CAR T cells were then activated under Th1 or Treg polarizing conditions in the absence or presence of αKG for 6-7 days and adoptively transferred into 24JK-ERB tumor-bearing mice. At day 20 post tumor inoculation, the percentages of adoptively transferred ErbB2-CAR CD4 T cells in draining lymph nodes (dLN) were not significantly different in any of the groups (**Figure 3B**). Tumor infiltration of ErbB2-CAR CD4 T cells was significantly higher for those lymphocytes activated under Th1 conditions as compared to Treg conditions, with relative intra-tumoral T cell percentages of 52±6% and 23±4%, respectively (p<0.001; **Figure 3C**). Notably though, T cell infiltration was markedly enhanced following *ex vivo* activation under Treg polarizing conditions in the presence of αKG, increasing to levels equivalent to those detected for Th1-polarized cells (50±2%, p=0.002 **Figure 3C**). Furthermore, the percentages of FoxP3^+^ ErbB2-CAR T cells in both LNs and tumors were significantly de creased by *ex vivo* activation under Treg polarizing conditions in the presence of αKG (p<0.001) and the potential of these cells to secrete IFNγ was augmented (p<0.01; **Figures 3B**). Thus, these data demonstrate that the *ex vivo* modification of αKG levels in adoptively transferred T cells can alter their *in vivo* intra-tumoral infiltration and persistence. Moreover, *ex vivo* αKG treatment significantly increased the potential of these tumor-specific T cells to secrete IFNγ *in vivo*. Thus, αKG is essential to maintain the proper Th1/Treg balance for *in vivo* anti-tumoral responses.

### Gene ontology analysis of αKG reprogrammed T cells reveals significant alterations in genes involved in lipid and mitochondrial pathways

To gain insights into the mechanisms via which αKG enhances Th1 differentiation while negatively regulating Treg differentiation, RNASeq analysis was performed. In accord with the alteration in T cell fate under Treg but not Th1 conditions, αKG-mediated changes in gene expression were more robust in Treg conditions (**Figure 4A**). There were substantially more downregulated as well as upregulated genes in Treg as compared to Th1 conditions (237 genes in Th1 conditions and 1,639 genes in Treg conditions with an adjusted p<0.01; **Supplemental Tables 1 and 2**). In Treg conditions, upregulation of transcription factors such as T-bet (*Tb×21*, padj=1.1×10^-56^) were expected based on our initial qRT-PCR analyses (**Figure 2C**) but other transcription factors such as Irf1, Irf8, and Stat1 were also highly upregulated (padj<5×10^-7^), suggesting increased signaling through interferon regulatory factors in these cells (**Figure 4B**). Interestingly though, housekeeping control genes were unaffected (Rpl11, Ube3c; **Figure 4B**), highlighting the specificity of this response. Most importantly, over-representation analysis of gene ontogeny (GO) terms for non-redundant biological processes (BP) using the WebGestalt program WebGestalt (Liao et al., 2019) revealed the importance of lipid and mitochondrial metabolism (**Figure 4C**). Within the 856 downregulated genes (protein-encoding), evaluation of non-redundant biological processes revealed the importance of lipid and phospholipid processes while mitochondrial processes were amongst the most highly upregulated biological processes (726 protein-encoding upregulated genes; **Figure 4C**). These data strongly highlight the importance of αKG-induced metabolic reprogramming in mediating cell surface remodelling under conditions of attenuated Treg polarization.

**Figure 4:**
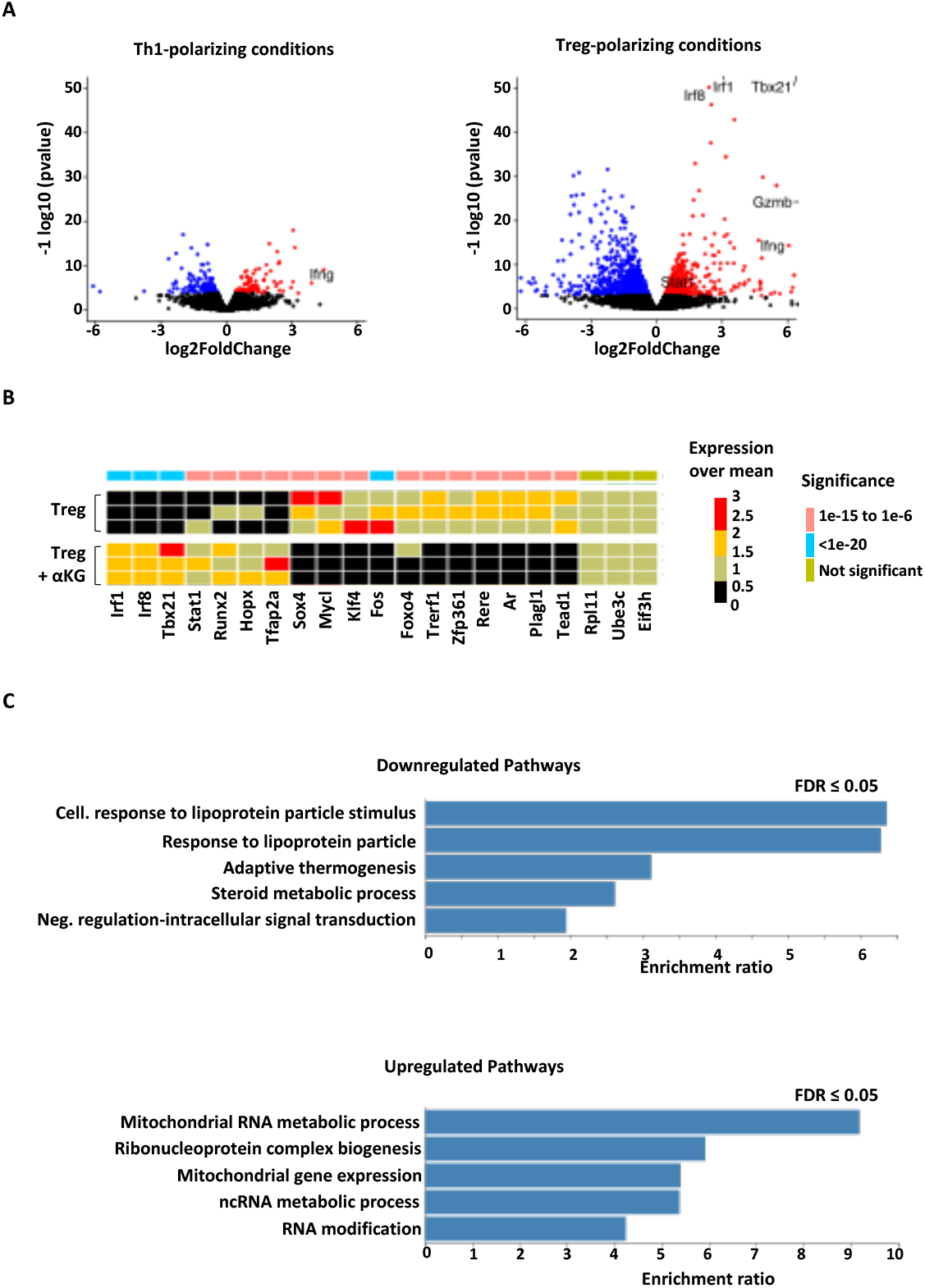
Genes affecting membrane-related processes are significantly impacted by αKG in Treg polarizing conditions. **(A)** Naïve CD4 T cells were activated under Th1 or Treg polarizing conditions in the presence or absence of exogenous αKG. On day 4 following polarization, cells were collected and transcripts analyzed by RNAseq. Volcano plot representations of the differential expression analyses of genes expressed in T cells activated in Th1- (left) and Treg-polarizing (right) conditions in the absence or presence of αKG are presented. Transcripts that are differentially expressed (FDR < 0.05 and baseMean expression > 100) are colored. **(B)** Presentation of alterations in transcription factor genes following activation in Treg polarizing conditions in the absence or presence of αKG. The level of statistical significance is indicated in the color scheme, as shown in the legend (right). **(C)** Overrepresentation analyses of Gene ontology (GO) for non-redundant biological processes within downregulated (top, 856) and upregulated (bottom, 726) genes following activation under Treg polarizing conditions in the presence of αKG were evaluated by WebGestalt {Liao, 2019 #344}. The different GO biological processes are presented on the y-axis and the enrichment ratio for each GO term is presented. Full data sets for downregulated and upregulated genes can be found at http://www.webgestalt.org/results/1597787239/# and http://www.webgestalt.org/results/1597787082/#, respectively.

### Lipid metabolism and lipidome remodelling are linked to αKG-mediated inhibition of Treg differentiation

Based on the massive changes in the transcription of genes associated with lipid processes and as “membranes” were in the top GO category of the cellular components analysis (367 of 824 categories), we hypothesized that αKG affected the composition, organization, and function of T cell membranes. As a first test of this hypothesis, we evaluated the impact of αKG on the lipid content of Treg-polarized CD4 T cells. Notably, increasing intracellular αKG resulted in a massive increase (~2-fold) in total cellular lipids (p<0.0001; **Figure 5A**). Because plasma membranes contain more than 90% of lipids species, we next wondered whether αKG altered membrane composition. We first focused on cholesterol, a hallmark of membrane dynamics (Needham and Nunn, 1990; Sezgin et al., 2017). We observed decreased levels of the ATP-binding cassette transporters ABCA1 and ABCG1 (**Supplemental Figure 3A**), which are responsible for cholesterol efflux (Yvan-Charvet et al., 2010). However, a similar relative abundance of cholesterol in the lipid membrane was observed in our lipidomic analyses in the absence or presence of αKG (**Figure 5B**). Moreover, cholesterol supplementation was not sufficient to rescue the αKG-mediated attenuation of Treg differentiation (**Supplemental Figure 3B**). Indeed, we found that cholesterol homeostasis in these cells, in the context of low levels of cholesterol efflux genes, was likely compensated by decreased levels of genes involved in cholesterol biosynthesis after αKG treatment (**Supplemental Figure 3C**). Several other major membrane lipid species, such as phosphatidylcholine (PC), phosphatidylinositol (PI) and phosphatidylserine (PS) were also not significantly altered in αKG-treated Treg (**Figure 5B**).

**Figure 5:**
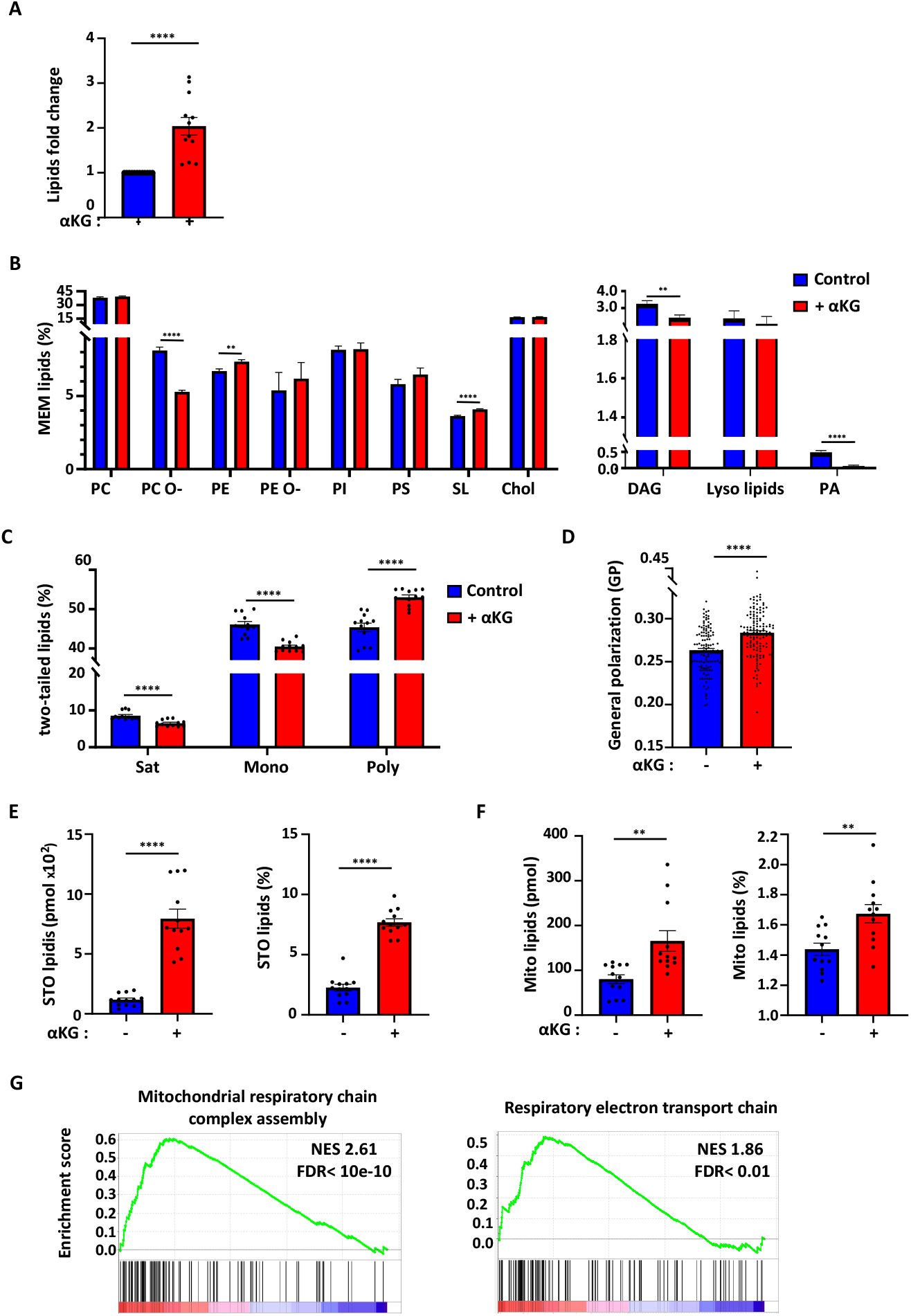
Alterations in lipidome remodeling in Treg-polarized cells in response to αKG are associated with dramatic increases in storage and mitochondrial lipids. **(A)** Lipidomic analysis of total lipid levels in CD4 T cells polarized to a Treg fate in the absence (blue) or presence (red) of αKG (control Treg conditions are arbitrarily presented as “1” and data represent 4 independent experiments performed in triplicate). Each point presents a sample with means ± SEM presented. (**B**) The abundance of the different membrane-lipid classes (PC, phosphatidylcholine; PC O-, PC plasmalogens; PE, phosphatidylethanolamine; PE O-, PE plasmalogens; PI, phosphatidylinositol; PS, phosphatidylserine; SL, sphingolipids; Chol, cholesterol; DAG, diglyceride; Lyso lipids (LPA, lysophosphatidic acid; LPC, lysophosphatidylcholines; LPE, lysophosphatidylethanolamine; LPC O-/ LPE O-, LPC and LPE plasmalogens); and PA, phosphatidic acid) was determined and means ± SEM are presented. (**C**) The level of unsaturation in twotailed membrane lipids were evaluated in the absence or presence of αKG (Sat, Saturated; Mono, Monounsaturated; Poly, Polyunsaturated) and means ± SEM are presented. **(D)** Quantification of membrane packing, presented as generalized polarization (GP), was evaluated following Treg polarization in the absence (129 cells) or presence of αKG (132 cells). Means ± SEM from 3 independent experiments are presented. **(E)** The total abundance (pmol; picomol) of storage lipids (TAGs, triglyceride and CEs, cholesterol esters) was evaluated (left) as well as their relative abundance as a function of total lipids (right) in the indicated conditions. Means ± SEM are presented. **(F)** The total abundance of mitochondrial lipids (CL, cardiolipin and PG, phosphatidylglycerol) was evaluated (left) and their abundance as a function of total lipids (right) are presented. (**G**) GSEA enrichment plots of mitochondrial respiratory chain complex assembly and respiratory electron transport chain are presented. The green curve corresponds to the enrichment score, and the barcode plot indicates the position of the genes in each gene set with the most upregulated genes presented in red and the most downregulated genes presented in blue. Statistical differences were determined by an unpaired 2-tailed t-test (*, p<0.05; **, p<0.01; ***, p<0.001; ****, p<0.0001).

In contrast, quantitative shotgun lipidomics revealed highly significant changes in the unsaturation state of phospholipids; decreased levels of saturated and mono-unsaturated lipids and conversely, increased polyunsaturated lipids (p<0.0001; **Figure 5C**). Consistent with these data, several desaturase enzymes including SCD1, SCD2, FADS1, and FADS2 were markedly impacted by αKG in Treg conditions (**Supplemental Figure 4A** and **Supplemental Table 1**). Interestingly, transcripts encoding these genes as well as ELOVL elongases were significantly altered by αKG in conditions of Treg but not Th1 differentiation (**Supplemental Figure 4A** and data not shown). Thus, αKG-mediated decreases in desaturase enzymes were coupled to a concordant perturbation in membrane lipid unsaturation under conditions of T cell reprogramming. Furthermore, other phospholipids were significantly altered by αKG-mediated T cell reprogramming; phosphatidylethanolamine (PE) and sphingolipids (SL) were augmented (p<0.01) and the alkyl-ether-linked phosphatidylcholine (PC O-), a plasmalogen whose orientation of the polar head group differs with respect to the membrane surface from diacyl phosphatidylcholine (Han and Gross, 1990), was decreased (p<0.0001; **Figure 5B**).

These robust lipidomic findings led us to evaluate changes in the physical properties of the membrane in the polarized CD4 T cells. Using the polarity Di-4-AN(F)EPPTEA probe and spectral imaging (Sezgin et al., 2015a; Sezgin et al., 2019), we determined that αKG treatment resulted in significantly increased membrane packing, quantified by the dimensionless parameter generalized polarization (GP), which reports on the fluorescence emission shift of the membrane sensitive fluorophore (p<0.0001, **Figure 5D** and **Supplemental Figure 3D**). This small yet significant change in GP is consistent with the broad lipid remodelling in response to αKG treatment. While incorporation of polyunsaturated fatty acids (PUFAs) into phospholipids generally increases fluidity (Ernst et al., 2018; Levental et al., 2016; Sezgin et al., 2015a; Sezgin et al., 2017), these data strongly suggest that complex compensatory changes between the lipidome-wide remodelling and high levels of PUFAs (as recently reported in (Levental et al., 2020) contribute to increased membrane packing and rigidity. Thus, αKG-mediated perturbation of lipid unsaturation and phospholipids during Treg differentiation leads to increased membrane packing and rigidity.

### αKG couples membrane lipid accumulation to fatty acid storage and mitochondrial lipid biosynthesis

Fatty acids are not only components of plasma membrane lipids but are also present in the cells, where they are stored as triacylglycerides (TAGs). Such TAGs are a major source of stored energy for the cell, presenting a dynamic pool of fatty acids that can be rapidly mobilized in response to cellular stress and energy requirements (Howie et al., 2019; Jarc and Petan, 2019). Notably, we found that αKG dramatically increased both the absolute levels of storage lipids, from 117±14 to 795±79 pmol/10e6 cells, representing an increase from 2.3±0.3% to 7.7±0.3% of total lipids (p<0.0001; **Figure 5E**). TAG metabolism has been associated with Treg function but our data show that increased storage lipids are associated with a decreased potential for Treg differentiation. Indeed, DGAT1, responsible for esterifying exogenous fatty acids into triglycerides (Bhatt-Wessel et al., 2018), was expressed at significantly higher levels in Th1 than Treg but was significantly upregulated by αKG, to levels similar to those expressed in Th1 cells (padj=0.001; **Supplemental Figure 4B** and data not shown). These data reveal the critical role of αKG in increasing the energy storage of CD4 T cells activated in Treg-polarizing conditions.

Increased membrane lipid composition with subsequent lipid storage in lipid droplet often parallels mitochondrial lipid biosynthesis (Jarc and Petan, 2020; Pernes et al., 2019). A close interaction between the presence of the two non-bilayer forming phospholipids in the mitochondrial inner membrane, PE and cardiolipin (CL) could reflect the close association between lipid metabolism and mitochondrial function (Schenkel and Bakovic, 2014). We found that αKG dramatically increased both CL and PG, quantified as mitochondrial lipids (Kojima et al., 2019; Pennington et al., 2019); the proportion of mitochondrial lipids increased from 80±10 to 166±23 pmol/10^6^ cells (**Figure 5F**, left panel), accounting for 1.4±0.04% and 1.7±0.06% of total membrane lipids, respectively (p<0.01; **Figure 5F**, right panel). Because CL biosynthesis is linked to efficient mitochondrial metabolism (Gohil et al., 2004), these data suggest a role for αKG-mediated lipid metabolism in promoting mitochondrial respiration. Indeed, GSEA revealed that “mitochondrial respiratory chain complex assembly” as well as “respiratory electron transport chain” were significantly enriched by αKG under conditions of Treg polarization (**Figure 5G**). Together, these results reveal the critical role of αKG in increasing mitochondrial-related energy storage in Treg-polarizing conditions.

### Augmented αKG-dependent TCA cycling and OXPHOS negatively regulates Treg differentiation

Based on the significant increase in the transcription of mitochondrial respiratory genes as well as mitochondrial lipid metabolism, we evaluated the contribution of αKG to mitochondrial respiration in Treg polarizing conditions. While mitochondrial metabolism is critical for Treg differentiation and function (Beier et al., 2015; Chapman et al., 2018; He et al., 2017; Howie et al., 2017; Layman et al., 2017; Weinberg et al., 2019), long chain fatty acid oxidation does not appear to be required (Raud et al., 2018) and the precise role of the TCA cycle in fueling OXPHOS during induced Treg generation is not known. Interestingly, the basal oxygen consumption rate (OCR, correlating with OXPHOS) was significantly lower in Treg-than Th1-polarizing conditions (p<0.01; **Figure 6A**). These data are in accord with a recent study that reported a lower OXPHOS and glycolysis in activated tTreg as well as TGFβ-induced Treg as compared to Th0 and Th1 polarized cells (Priyadharshini et al., 2018). The lower OXPHOS in Treg cells (as compared to Th1) was associated with an augmented glycolysis, measured as a function of the extracellular acidification rate (ECAR, **Figure 6A**). Thus, by day 4 of polarization, CD4 T cells polarized to a Treg fate exhibit a low energetic profile as compared to cells polarized to a Th1 fate.

**Figure 6:**
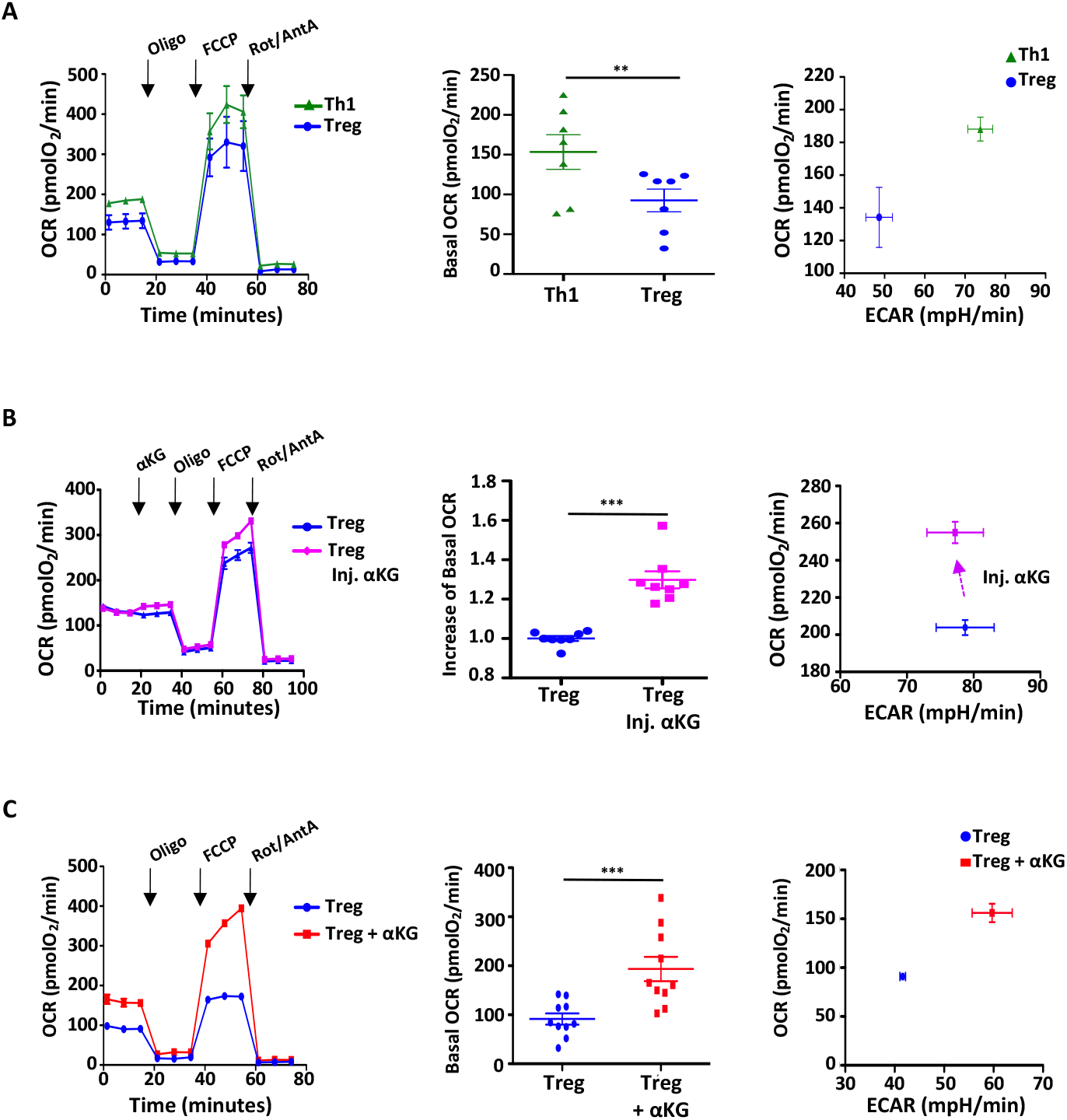
Lipid remodeling in response to αKG-treatment is associated with an augmentation in oxidative phosphorylation. **(A)** Oxygen consumption rate (OCR), a measure of oxidative phosphorylation, was monitored at day 4 of Th1 and Treg polarization on a Seahorse XFe96 analyzer following sequential injection of oligomycin, FCCP and antimycin A/rotenone (arrows; left panel). Mean basal OCR levels ± SEM are presented (middle panel, n=7 independent experiments). Energy plots of basal OCR (y axis) and extracellular acidification rate (ECAR) (x axis), a measure of glycolysis, are presented (right). **(B)** The immediate impact of αKG on cell metabolism was evaluated by injecting αKG directly into the XFe96 flux analyzer at day 2 of Treg polarization (left panel; data are representative of 1 of 8 independent experiments). Quantification of the fold change in basal OCR (middle panel) and OCR/ECAR energy plots (right) are presented. **(C)** OCR of CD4 T cells activated for 4 days under Treg-polarizing conditions in the absence or presence of αKG, are presented (left panel; data are; representative of 1 of 10 independent experiments). Quantification of the mean OCR ± SEM in 10 independent experiments are shown in the middle panel and OCR/ECAR energy plots are presented on the right. Significance was determined by a 2-tailed paired t-test (*, p<0.05; **, p<0.01; ***, p<0.001; ****, p<0.0001; NS, not significant).

We found that αKG directly affected the metabolism of Treg-polarized cells because its injection into the flux analyzer immediately increased oxygen consumption and the OCR/ECAR ratio (p<0.001; **Figure 6B**). Furthermore, this impact of αKG was maintained, with significant increases in both basal and maximal respiratory capacities following activation in Treg-polarizing conditions for 4 days (p<0.001; **Figure 6C**). Notably, the αKG-mediated augmentation in the OCR/ECAR energetic profile was coupled to a significantly increased ATP/ADP ratio (p<0.05; **Figure 7A**). These data strongly suggested that αKG increased cycling of TCA intermediates (**Supplemental Figure 5A**). Indeed, αKG levels were markedly elevated (p<0.0001), as expected from the ectopic addition of mM quantities, as were malate levels (p<0.05; **Figure 7B**). Furthermore, we detected a substantial utilization of ectopic αKG in the TCA cycle, monitored by the tracing of glutamine and glucose uniformly labelled with heavy carbon (^13^C_5_ and ^13^C_6_, respectively)—in the presence of αKG, there was a significantly increased contribution of non-labelled glucose or glutamine carbons (C_0_) in the generation of the different intermediates (**Supplemental Figure 5B**). The enhanced level of TCA cycling was also associated with a decreased generation of intracellular lactate and glutamate (**Supplemental Figure 5C**), highlighting the altered metabolism of these cells.

**Figure 7:**
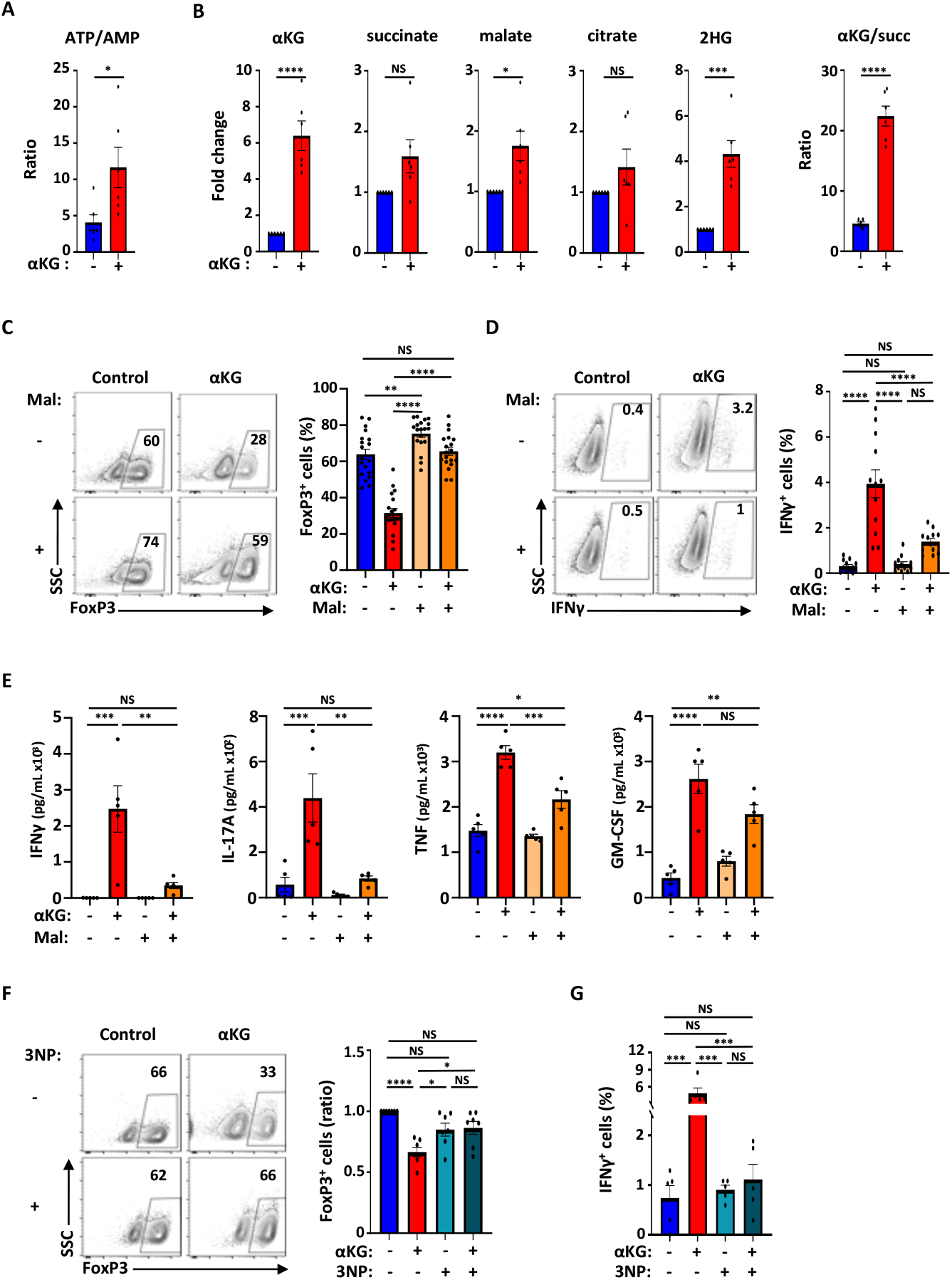
The αKG-mediated inhibition of Treg differentiation is attenuated by mitochondrial complex II inhibitors. **(A)** The ATP/AMP ratio in CD4 T cells activated in Treg polarizing conditions in the absence or presence of αKG were evaluated by mass spectrometry. Means ± SEM are presented (technical triplicates from 2 independent experiments). **(B)** The levels of αKG, succinate, malate, citrate and 2-hydroxyglutarate (2HG) were evaluated by mass spectrometry and the foldchanges in αKG-treated cells as compared to control conditions (set at “1”) are presented. The ratio of αKG to succinate in the different conditions is presented on the right. **(C)** The impact of malonate (Mal, 10 mM) on Treg polarization was evaluated in the absence or presence of αKG. Representative dot plots of FoxP3 expression are shown at day 4 (left) and quantification of the means ± SEM are presented (right, n=19). **(D)** Representative dot plots showing the percentages of IFNγ^+^ cells (left) as well as a quantification of the means ± SEM are presented at day 4 (right, n=12). **(E)** IFNγ, IL-17A, TNF, and GM-CSF secretion from CD4 T cells activated in Treg-polarizing conditions in the absence or presence of αKG and/or Mal was evaluated at day 3 and means ± SEM are presented (n=5). **(F)** The impact of 3-NP (62.5μM) on Treg polarization was evaluated in the absence or presence of αKG. Representative FACS plots of FoxP3 expression are presented at day 4 (left) and quantification of the means ± SEM are presented (right, n=7). **(G)** The percentages of IFNγ^+^ cells in the different conditions are presented as means ± SEM (n=5). Significance was determined either by unpaired 2-tailed test (panels A and B) or by a one-way ANOVA and Tukey multiple comparison test (panels C-G; *, p<0.05; **, p<0.01; ***, p<0.001; ****, p<0.0001; NS, not significant).

As conditions that alter the αKG/succinate ratio have been shown to regulate the balance between stem cell self-renewal and differentiation (Carey et al., 2015; TeSlaa et al., 2016) and we found that inhibition of Treg differentiation was associated with a significantly augmented αKG/succinate ratio (**Figure 7B**), we monitored the impact of ectopic succinate (Succ) on differentiation. Addition of succinate partially attenuated the negative impact of αKG, increasing the percentages of FoxP3^+^ cells from 27±4% and 38±5% (**Supplemental Figure 6**). While this difference was not significant, succinate can function by inhibiting prolyl hydroxylase (PHD), thereby leading to stabilization of the HIF-1α transcription factor, and/or by decreasing TCA flux through the αKG dehydrogenase complex (King et al., 2006; Koivunen et al., 2007; Selak et al., 2005). To distinguish between potential cytosolic and mitochondrial effects of succinate, we specifically inhibited succinate oxidation with dimethyl malonate (Mal), impairing the oxidation of succinate to fumarate without altering PHD activity in the cytoplasm (Mills et al., 2016). Notably, under these conditions, malonate not only restored the differentiation of FoxP3^+^ T cells in the presence of αKG (p<0.0001) but also increased FoxP3 levels under control conditions (control, 64±3%; αKG, 32±2%; Mal + αKG, 66±2%;, Mal, 75±2%; p<0.01, **Figure 7C**). Furthermore, in the presence of αKG, malonate massively reduced the production of inflammatory cytokines including IFNγ, IL-17A, and TNF (p<0.01; **Figure 7D and 7E**).

To corroborate the potential of malonate to rescue the αKG-mediated inhibition of Treg differentiation, we tested the capacity of another mitochondrial-respiratory complex II inhibitor, 3-nitropropionic acid (3-NP). Like malonate, 3-NP significantly increased the differentiation of FoxP3^+^ T cells in the presence of αKG (p<0.05; **Figure 7F**) and massively reduced the percentage of cells secreting IFNγ (p<0.001; **Figure 7G**). Together with our data showing that αKG significantly enriched for respiratory electron transport genes (**Figure 5G**), these results indicate that maintaining a low basal level of TCA flux is central to Treg differentiation and the preservation of a non-inflammatory state.

## Discussion

The proliferation and differentiation of T lymphocytes is regulated by antigen and cytokine signals but the induction of metabolic pathways is also required, resulting in an integration of environmental cues in cell fate decisions. Glutamine availability is a key determinant of T cell differentiation; CD4 lymphocytes undergo differentiation to IFNγ-secreting Th1 effectors under conditions of optimal glutaminolysis but are converted to FoxP3+ regulatory T cells when glutamine catabolism is abrogated (Klysz et al., 2015; Metzler et al., 2016; Nakaya et al., 2014). Here, we show that the pleiotrophic αKG metabolite, produced anaplerotically from glutamate, acts as a cell intrinsic check-point in CD4 T cell differentiation; αKG dramatically attenuated Treg generation while significantly augmented IFNγ secretion under both Th1- and Treg-polarizing conditions. Moreover, under conditions of Treg polarization, αKG induced the secretion of inflammatory cytokines that are associated with pathological Tregs. The potential of αKG to serve as an environmental cue that reprograms Treg differentiation was also observed in ErBB2-directed CAR-T cells, resulting in a significantly increased tumor infiltration and *in vivo* IFNγ secretion in ErbB2^+^ tumor-bearing mice. Using transcriptomic, metabolomic and lipidomic approaches, our study identifies a novel connection between αKG-mediated lipidome remodelling and mitochondrial reprogramming in attenuating polarization of naïve CD4 T lymphocytes to a Treg fate. Furthermore, we find that mitochondrial complex II, recently identified as an enforcer of terminal Th1 differentiation (Bailis et al., 2019), serves as a negative integrator of Treg differentiation.

T cell metabolism has elegantly been shown to integrate transcriptional profiles with epigenetic changes to modulate the differentiation state of the cell (Geltink et al., 2018; Patel and Powell, 2017; Yong et al., 2017a). Indeed, αKG specifically alters the epigenetic state of CD4 T cells by targeting DNA and histone methylation (Chisolm et al., 2017; Xu et al., 2017). This effect is modulated at least in part by αKG’s role as a required cofactor in the demethylase activity of TET dioxygenases (Lio and Rao, 2019; Rose et al., 2011; Tahiliani et al., 2009). Interestingly though, decreasing the conversion of glutamate to αKG was found to enhance Treg differentiation, and this effect was associated with a diminished reduction of αKG to the competitive 2-HG TET inhibitor (Xu et al., 2017). Similarly, we found that high intracellular αKG levels were associated with a significant increase in intracellular 2-HG levels (**Figure 7A**, p<0.0001). However, while αKG-dependent TET activity is required for the demethylation of the FoxP3 locus and its subsequent expression (Nakatsukasa et al., 2019; Yang et al., 2015; Yue et al., 2019; Yue et al., 2016), it is not clear whether 2-HG accumulation is responsible for the decreased Treg polarization detected in these conditions. The MS studies performed here show that even though 2-HG is markedly increased by ectopic αKG, the αKG/2-HG ratio is significantly higher than in the control conditions (1.4±0.06 to 2.1±0.1, p=0.0009). As FoxP3 expression was strongly induced in control Treg polarizing conditions but not in the presence of ectopic αKG, these data indicate that the altered differentiation state is not due to increased levels of 2-HG.

While αKG/2-HG ratios can alter the epigenetic state of a cell, the intracellular αKG/succinate ratio also contributes to cell fate via changes in histone and DNA demethylation (Carey et al., 2015; Liu et al., 2017; TeSlaa et al., 2016; Tischler et al., 2019) as does histone acetylation via the glycolytic generation of acetyl-coA (Moussaieff et al., 2015; Peng et al., 2016). Furthermore, increasing succinate levels promotes a pro-inflammatory macrophage phenotype by increasing mitochondrial reactive oxygen species (Mills et al., 2016). However, in contrast to the present study, previous research evaluating the impact of αKG/succinate ratios on cell fate found that its effect was independent of mitochondrial metabolism. Rather, the αKG/succinate ratio was shown to regulate differentiation through changes in histone and DNA demethylation (TeSlaa et al., 2016). That being said, metabolic alterations have been determined to reprogram T cell differentiation via other mechanisms including changes in binding of the CCCTC-binding factor (Chisolm et al., 2017), alterations in GAPDH activity (Balmer et al., 2016; Chang et al., 2013; Peng et al., 2016), and mTORC pathways (Johnson et al., 2018), amongst others.

In our study wherein αKG/2-HG and αKG/succinate ratios were significantly increased, our RNASeq data highlighted cholesterol biosynthesis as the metabolic pathway most impacted by αKG following activation of CD4 T cells under Treg polarization conditions. Key cholesterol biosynthesis genes such as Cyp51 were massively decreased (FDR=6×10^-5^) as were cholesterol exporters such as ABCA1 and ABCG1 (FDR<1×10^-6^). It was therefore surprising that our experiments altering cholesterol levels did not change Treg polarization efficiency but this may be due to the inability of supplemental cholesterol to compensate the specific intracellular membranes that were altered by αKG (e.g. plasma membrane, ER, etc). Nonetheless, plasma αKG levels are known to be modulated by the metabolic state of the organism: αKG levels are significantly increased in starved bacteria/yeast (Brauer et al., 2006) and moreover, high αKG levels inhibit ATP synthase and TOR, resulting in an augmented lifespan in worms (Chin et al., 2014). While one study reported an association between αKG levels, obesity and non-alcoholic fatty liver disease (Rodriguez-Gallego et al., 2015), most vertebrate studies are in accord with the non-vertebrate data, showing a protective role for αKG in modulating cholesterol and fatty acid metabolism. In response to acute exercise, serum αKG concentrations increase (Brugnara et al., 2012) and αKG promotes the absorption of fatty acids in muscle tissues of pigs (Chen et al., 2019). In accord with these results, αKG prevents hypercholesterolemia and weight gain in mice and rats fed with a high-fat diet (Nagaoka et al., 2020; Radzki et al., 2009) as well as in piglets after prenatal exposure to dexamethasone (Sliwa et al., 2009).

The data presented above strongly support a coupling of αKG to lipid metabolism. However, our study is the first, of which we are aware, to demonstrate a significant impact of αKG on the transcriptional profile of genes involved in the cholesterol pathway as well as more broadly, in membrane-associated genes. Although extensive research has focused on the contributions of fatty acid oxidation to lipid metabolism and cell fate decisions, there has been a paucity of research on the role of nonoxidative lipid pathways in regulating hematopoietic cell fate (reviewed in Lee et al., 2018; Pernes et al., 2019). Much of the lipid metabolism research in immune cells has focused on the incorporation of PUFAs into the membrane, with the ratio of n-3 to n-6 PUFAs found to modulate inflammation and autoimmunity (Kim et al., 2018; Monk et al., 2012; Shaikh and Edidin, 2006). While the role of lipidome remodelling in T lymphocyte fate has not yet been evaluated, recent studies have highlighted the importance of the cell’s lipidome in mesenchymal stem cell differentiation (Levental et al., 2017), myelopoiesis (Mitroulis et al., 2018), stem cell pluripotency (Wu et al., 2019), as well as extended lifespan (Schmeisser et al., 2019). In the study presented here, we find that the αKG-mediated attenuation of Treg polarization is coupled to a robust lipidomic remodelling.

In the context of the αKG-mediated phospholipid remodelling that was associated with a markedly attenuated Treg differentiation, we detected a >5-fold increase in TAGs, or storage lipids. These storage lipids, representing a highly reduced form of carbon, serve as an energy reserve that allow cells to meet increased cellular demands (reviewed in (Aon et al., 2014)). In accord with the impact of TAGs on increased energy reserves, ectopic αKG was associated with a >3-fold increase in the ATP/AMP ratio. Furthermore, TCA cycle intermediates were augmented and our metabolic tracing experiments highlighted the importance of carbons from ectopic αKG, replacing carbons from glucose or glutamine, in the generation of these intermediates. While Tregs have been found to use fatty acids for OXPHOS and lipid storage (Howie et al., 2017; Howie et al., 2019; Michalek et al., 2011), our data show that an αKG-mediated increase in DGAT1 (FDR=0.001), a rate-limiting enzyme in TAG accumulation and cholesterol absorption (Cases et al., 1998; Sachdev et al., 2016), as well as a concomitant increase in TAGs, were associated with a significantly diminished Treg differentiation potential.

The αKG-mediated lipidome remodelling that resulted in increased TAGs also led to a 2-fold increase in mitochondrial phospholipids (CL and PG, p<0.01), strongly suggesting that mitochondrial metabolism would be augmented in these conditions. Previous studies on the relative levels of OXPHOS in Tregs as compared to Teff have yielded discrepant results, likely due to differences in the study systems and the distinct types of Treg subsets that were being evaluated (Koch et al., 2009; Sun et al., 2018). Tregs have been found to exhibit increased OXPHOS as compared to Teff subsets (Angelin et al., 2017; Gerriets et al., 2016; Gerriets et al., 2015; Michalek et al., 2011) but within tumors, Tregs exhibit a gene signature consistent with glycolysis (Pacella et al., 2018). Furthermore, while thymic Treg (tTreg) function was found to be dependent on mitochondrial complex III function (Koch et al., 2009), the level of OXPHOS in induced Tregs was reported to be significantly lower than in either tTreg or Th1 cells (Priyadharshini et al., 2018). In accord with the latter study, we detected significantly lower levels of OXPHOS following activation under Treg-polarizing than Th1-polarizing conditions. Furthermore, αKG directly augmenting OXPHOS. However, the association of OXPHOS to Treg function is complex; inhibition of OXPHOS can suppresses Treg suppressive functions (Beier et al., 2015; Weinberg et al., 2019), but decreased mitochondrial integrity, due to loss of the fatty acid binding protein FABP5, has been found to enhance Treg function (Field et al., 2019). Here, we identified mitochondrial complex II as a critical negative regulator of Treg differentiation: abrogation of complex II by either malonate or 3-NP restored Treg polarization in the presence of αKG and decreased the production of inflammatory cytokines. In this regard, it is notable that abrogation of this complex was recently found to suppress terminal Th1 effector function (Bailis et al., 2019). Together, these data reveal an inverse relationship in ETC programs in specifying Treg vs Th1 effector differentiation.

The ability of the αKG metabolite to alter the fate of an activated CD4 T lymphocyte can have important clinical implications. As indicated above, the metabolic environment can alter *in vivo* serum αKG concentrations, as a function of starvation/obesity as well as exercise (Brauer et al., 2006; Brugnara et al., 2012; Rodriguez-Gallego et al., 2015; Zdzisinska et al., 2017). As such, αKG can potentially alter the activity of endogenous or adoptively-transferred tumor-specific T lymphocytes. Indeed, CAR-T cell metabolism, modulated as a function of the CAR co-stimulatory domain, has been shown to alter anti-tumor function (Kawalekar et al., 2016; Xu et al., 2019) and here, we find that short-term *ex vivo* αKG treatment enhances the *in vivo* persistence of CAR-T cells in a murine tumor model. αKG increased intra-tumoral infiltration of Treg-polarized CAR-T cells and moreover, was associated with levels of IFNγ secretion that were similar to those induced by Th1-polarized CAR-T cells. Furthermore, in the context of translational applications, we found that very similarly to its impact on murine T cell differentiation, αKG also significantly decreased human Treg differentiation (**Supplemental Figure 7**). Altogether, our study identifies the pleiotropic αKG metabolite as an intrinsic checkpoint regulator in T cell function, increasing IFNγ secretion by CD4 T cells as well as CD4 CAR-T cells, irrespective of whether they are activated in Treg or Th1 conditions. Furthermore, our findings reveal the coordinated regulation of CD4 T cell differentiation by metabolic modules that couple lipidome remodelling to mitochondrial reprogramming and cell fate.

## Materials and Methods

### Mice

Ly5.1, Ly5.2 C57BL/6 or transgenic mice harboring an ErbB2-CAR on all hematopoietic cells downstream of the Vav promoter as described (Yong et al., 2015) were crossed with FoxP3-GFP reporter mice (Wang et al., 2008). Mice were housed in a conventional, pathogen-free facility at the Institut de Génétique Moléculaire de Montpellier. Animal care and experiments were approved by the local animal facility institutional review board in accordance with French national guidelines.

### T cell isolation, activation and culture

Murine CD4^+^ T cells were purified using the MACS CD4^+^ T cell negative selection kit (Miltenyi Biotec) and naïve CD4^+^ T cells from WT, FoxP3-GFP and ErbB2-CAR transgenic mice were then sorted on the basis of a CD4^+^CD8^-^CD62L^+^CD44^-^GFP^-^CD25^-^ expression profile on a FACSAria flow cytometer (BD Biosciences). Thymic Tregs (tTregs) were sorted from FoxP3-GFP reporter mice based on a CD4^+^CD8^-^CD62L^+^CD44^-^GFP^+^ profile. T cell activation was performed using plate-bound α-CD3 (clone 145-2C11, 1 μg/ml) and α-CD28 (clone PV-1, 1 μg/ml) monoclonal antibodies in RPMI 1640 medium (Life Technologies) supplemented with 10% FCS, 1% penicillin/streptomycin (Gibco-Life technologies) and β-mercaptoethanol (50 μM). For Th1 and Treg-polarizing conditions, IL-12 (15 ng/ml) and α-IL-4 antibody (5 ug/ml) or hTGF-β (3 ng/ml) and rhIL-2 (100 U/ml), respectively, were added to the cultures. When indicated, cell permeable αKG (dimethyl ketoglutarate, 3.5 mM; Sigma), malonate (dimethyl malonate, 10mM; Sigma), succinate (diethyl succinate, 3.5 mM; Sigma), 3-NP (3-Nitropropionic acid, 62.5 μM; Sigma) and water-soluble cholesterol (50 mM, Sigma) were added. Cells were split 3 days later with media supplemented with rhIL-2 (100U/mL) and drugs were added at their original concentration. Cells were maintained in a standard tissue culture incubator containing atmospheric O_2_ and 5% CO_2_.

Human CD4^+^ T cells were isolated from healthy adult donors. All experiments using primary human cells were conducted in accordance with the Declaration of Helsinki and IRB approval to the French Blood Bank (Etablissement Francais du Sang). T lymphocytes were purified by negative-selection using Rosette tetramers (Stemcell Technologies Inc) and the purity was monitored by flow cytometry. For sorting of human CD4^+^ naïve T cells, CD4^+^ T cells were sorted on the basis of a CD4^+^CD8^-^CD45RA^+^CD45RO^-^CCR7^+^CD62L^+^CD127^+^CD25^-^ phenotype on a BD FACSAria flow cytometer. For Treg-polarizing conditions, hTGF-β (3 ng/ml) and rhIL-2 (100 U/ml) were added to the cultures. T cell activation was performed using plate-bound α-CD3 (clone OKT3, Biolegend) and α-CD28 (clone 9.3) mAbs at a concentration of 1 μg/ml in RPMI medium 1640 (Life Technologies) supplemented with 10% FCS and 1% penicillin/streptomycin (Gibco, ThermoFisher).

### Immunophenotyping and flow cytometric analysis

Immunophenotyping of cells was performed with fluorochrome-conjugated antibodies, and intracellular staining was performed after the fixation and permeabilization of the cells (intracellular staining kit from ThermoFisher or BD Biosciences). Analysis of cells collected from *in vivo* experiments were performed following Fc block (2.4G2, BioXcell) for 10 minutes prior to incubation with conjugated antibodies for FACS analysis. For intracellular cytokine staining, cells were stimulated with phorbol 12-myristate 13-acetate (PMA) (100 ng/ml) and ionomycin (1 μg/ml) in the presence of brefeldin A (10 μg/ml; all from Sigma) for 3.5–4h at 37°C. Cells were labelled with fixable viability dye prior to fixation, followed by at least 3 hours of intracellular staining with the following antibodies; FoxP3-PeCy7, T-Bet-PE and IFNγ-APC. Cytokine production (IFNγ, TNF, IL-17A, GM-CSF) was also assessed by Cytometric Bead Array (CBA) Kit (BD Biosciences) at day 3 of polarization. Cells were analyzed on a FACSCanto or LSRII-Fortessa flow cytometer (BD Biosciences). Data analysis was performed using FlowJo (Tree Star software) and FCAPArray Software (CBA analysis). Murine and human antibodies are listed in **Supplemental Tables 3 and 4**, respectively.

### Tumor model and adoptive T cell transfer

The murine 24JK fibrosarcoma cell line expressing the human ErbB2/Her2 antigen was generated as described (Yong et al., 2016). 24JK-ERB cells (1×10^6^) were subcutaneously injected into RAG2^-/-^ mice and allowed to establish over a 7-day period. On day 7, 3-5 x 10^6^ *in vitro* ErbB2-CAR T cells, activated as indicated, were injected retro-orbitally into tumor-bearing mice. For endpoint analyses, tumors were excised, mechanically digested in T cell culture media (as above) using collagenase IV (1 mg/ml, Sigma) and DNASE I (500μg/ml, Roche), incubated for 25 minutes at 37°C and processed into single cell suspensions for FACS analysis. Lymph nodes and spleens of tumor bearing mice were isolated and dissociated into single cell suspensions in PBS + 2% FCS and analyzed by flow cytometry.

### RNAseq analysis

T cells (5×10^6^) from triplicates of 2 independent experiments were lysed in TRIzol reagent (ThermoFisher). Total RNA was isolated per manufacturer’s instruction and resuspended in RNase free water. RNA was quantified with Qubit^®^ RNA HS Assay Kit (Thermo Fisher Scientific) and RNA intergrity was verified with Agilent RNA 6000 Nano kit on the 2100 Bioanalyzer (Agilent Technologies) according to the manufacturers’ protocol. RNA-Seq libraries were prepared from 65 ng of total RNA with ERCC ExFold RNA Spike-In Mix 1 (Life Technologies) of 1 μL of 1:5000 (v/v) dilution with TruSeq^®^ Stranded mRNA Library Prep for NeoPrepTM kit as per the NeoPrep Library Preparation System Guide on the instrument (Illumina). The libraries were diluted 1:5 (v/v), with 1 μL used to check the expected size (~300 bp) and the samples’ purity were verified with the Agilent High Sensitivity DNA Kit as manufacturer’s recommendation (Agilent Technologies). One μL of the diluted libraries were used to quantify as recommended in the Quant-iT PicoGreen dsDNA Assay Kit (Thermo Fisher Scientific) with standard range from 0 to 14 ng/ well. Pooled libraries of 1 nM were denatured and diluted as per the Standard Normalization Method for the NextSeq^®^ System (Illumina). Final loading concentration of 1.7 pM and 1% (v/v) PhiX was used with the NextSeq^®^ 500/550 High Output Kit v2 (150 cycles) (Illumina Samples were sequenced paired-end with 75 cycles of each read and single index of 6 cycles.

Reads were aligned to mm10 by STAR (version 2.5.2a) (Dobin et al., 2013). NCBI’s genome-build GRCm38.p5 (12-2016) was used as the gene model. ERCC spike-in controls were added for normalization. The alignments to transcriptome were quantified using RSEM (version 1.2.31) (Li and Dewey, 2011). Differential expression was evaluated using the DESeq2 package (Love et al., 2014) and analyzed using the WebGestalt (WEB-based Gene SeT AnaLysis Toolkit) functional enrichment analysis web tool (Liao et al., 2019).

### Gene expression analysis

RNA was isolated from T cells as indicated using the RNeasy Micro Kit and then reverse-transcribed to produce cDNA using oligonucleotide priming with the QuantiTect Reverse Transcription Kit (both Qiagen). Quantitative PCR was performed using the QuantiTect SYBR green PCR Master mix (Roche) with 10 ng of DNA and 0.5 μM primers in a final volume of 10 μl. Amplification of DNA was performed using the LightCycler 480 (Roche). Each sample was amplified in triplicate. Relative expression was calculated by normalization to HPRT and primer sequences are presented in **Supplemental Table 5**.

### Lipid analyses

Mass spectrometry-based lipid analysis was performed at Lipotype GmbH (Dresden, Germany) as previously described (Sampaio et al., 2011) and triplicate samples (1×10^6^) were evaluated. Briefly, lipids were extracted using a two-step chloroform/methanol procedure (Ejsing et al., 2009). Samples were analyzed by direct infusion on a QExactive mass spectrometer (Thermo Scientific) equipped with a TriVersa NanoMate ion source (Advion Biosciences) as described (Mitroulis et al., 2018).

Data were analyzed as previously described, using an in-house lipid identification software based on LipidXplorer (Levental et al., 2020; Wang and Han, 2016). Analyses were only performed on samples with a signal-to-noise ratio >5 and a 5-fold higher signal intensity than in corresponding blank samples. Lipidomic analysis yielded a list of >600 individual lipid species. Quantities were determined by first transforming all detected lipids into mol% as a function of total lipids. Molar percentages of membrane lipids were determined after removing TAG and CE from the analysis. Storage lipids were analyzed as a function of TAGs and CE. Class composition was further evaluated in the datasets as well as the number of carbons and level of unsaturation in individual species.

### Spectral imaging and generalized polarization

Spectral imaging was performed as previously described (Levental and Levental, 2015; Levental et al., 2016; Levental et al., 2017; Sezgin et al., 2012; Sezgin et al., 2015b). Following staining of cells with 1 μg/ml Di-4-AN(F)EPPTEA for 5 min at room temperature, cells were evaluated by confocal based spectral imaging (Sezgin et al., 2015b). Di-4-AN(F)EPPTEA was excited with 488 nm laser and emissions were collected with a spectral channel in the spectral region 410-691 nm. For GP calculations, intensities at 560 nm and 650 nm wavelengths were used as ordered and disordered signal (*I_560_* and *I_650_*), respectively. GP was calculated as:

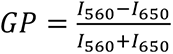

GP analysis was performed with custom GP plugin for ImageJ (Sezgin et al., 2015b).

### Mass spectrometry (LC-MS)

For profiling of intracellular metabolites, cells (1×10^6^) were treated as previously described (Oburoglu et al., 2014). Briefly, cells were incubated in RPMI medium without glucose or glutamine and supplemented with uniformly labelled [U-^13^C_6_]glucose or [U-^13^C_5_-^15^N_2_]glutamine, respectively, for 24 hours. Cells were rapidly washed in ice-cold PBS and metabolites were extracted in a 50% methanol / 30% acetonitrile/ 20% water solution. The volume of extraction solution was adjusted to 2×10^6^ cells/ml for all samples and were normalized to the number of cells obtained at the end of the 24-hour incubation period. All cultures were performed in triplicate. Both cell extracts and media were centrifuged at 16,000g for 10min at 4°C and supernatants were analyzed by HPLC-MS. Samples were separated on a Thermo Scientific Accela liquid chromatography system coupled with a guard column (Hichrom) and a ZIC-pHILIC column (Merck). Separations were performed using a gradient elution in a solution of 20mM ammonium carbonate, with 0.1% ammonium hydroxide and acetonitrile. Metabolites were detected using a Thermo Scientific Exactive Orbitrap mass spectrometer with electrospray ionization, operating in a polarity switching mode. Raw data were analyzed with Xcalibur 2.2 and LCquan 2.7 Thermo scientific software.

### Extracellular flux analysis

OCR and ECAR were measured using the XFe96 Extracellular Flux Analyzer (Seahorse Biosciences, Agilent). Cells (1.7-2×10^5^/well) were resuspended in XF media (buffered RPMI) in the presence of glucose (11 mM) and L-glutamine (2 mM), added to poly-D-lysine-coated wells (0.1mg/ml, Millipore) and monitored in basal conditions and in response to oligomycin (1 μM), FCCP (1 μM), rotenone (100 nM) and antimycin A (1 μM; Sigma). Metabolites were added as indicated. Basal respiration is calculated as followed: baseline cellular OCR-Non-mitochondrial respiration (post injection of Antimycin A/Rotenone).

### Statistical analyses

Data were analyzed using GraphPad software version 8 (Graph Pad Prism, La Jolla, CA) and p values were calculated using a one-way ANOVA (Tukey’s post-hoc test), unpaired or paired 2-tailed t-tests as indicated.

## Acknowledgments

We thank all members of our laboratories for discussions and scientific critique. We are indebted to Ünal Coskun, Michal Grzybek, Alessandra Palladini and Triantafyllos Chavakis for all their expertise and assistance with lipidomics analyses. We are grateful to Myriam Boyer and Stéphanie Viala of Montpellier Rio Imaging for support in cytometry experiments. We are very grateful to Amal Makrini of the SERANAD bioinformatics platform for her assistance with analyses. MIM was funded by the French Ministry of Health and a fellowship from ARC and CSY by an Australian doctoral fellowship and ARC. S.T. is supported by funding from Cancer Research UK (C596/A17196, Award 23982). This research was in part supported by the NIH Intramural Research Program of NIAID (S.A.M.) and NCI (N.T.). This work was supported by generous funding from the French national (ANR) research grants (NutriDiff and PolarAttack), FRM, ARC, Sidaction, ANRS, and the French laboratory consortiums (Labex) EpiGenMed and GR-EX.

