## Supplemental Figures for "αKG inhibits Regulatory T cell differentiation by coupling lipidome remodelling to mitochondrial metabolism"

**Supplemental Figure 1: Treg polarization is attenuated by αKG.**

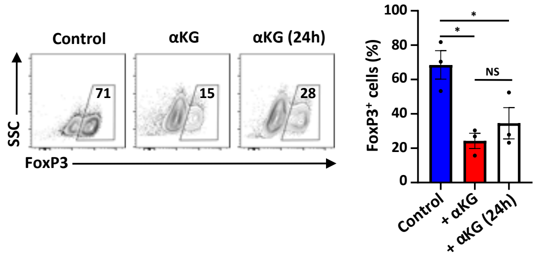


Naïve CD4 T cells were activated under Treg-polarizing conditions and αKG was added either at the time of activation or at 24h post stimulation. FoxP3 expression was evaluated at day 4 and representative dot plots are shown (left) as well as a quantification of the means ± SEM of 3 independent experiments (right). Significance was assessed by a one-way ANOVA and Tukey multiple comparison test (*, $p<0.05$; NS, not significant).

**Supplemental Figure 2: αKG inhibits Treg polarization of ErbB2-CAR T cells.**

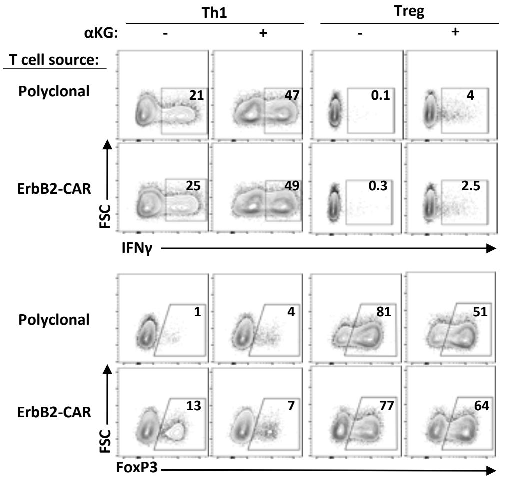


**(A)** The potential of transgenic ErbB2-CAR T cells, as compared to polyclonal T cells, to be polarized to a Th1 or Treg fate was compared in the absence or presence of αKG. IFNγ production was evaluated at day 4 of stimulation in the indicated conditions and representative plots are shown (top plots). The induction of FoxP3 expression was also evaluated at day 4 and the percentages of FoxP3$^{+}$ cells are indicated (bottom plots).

**Supplemental Figure 3: Decreased αKG-induced transcription of cholesterol-biosynthesis genes does not regulate Treg polarization.**

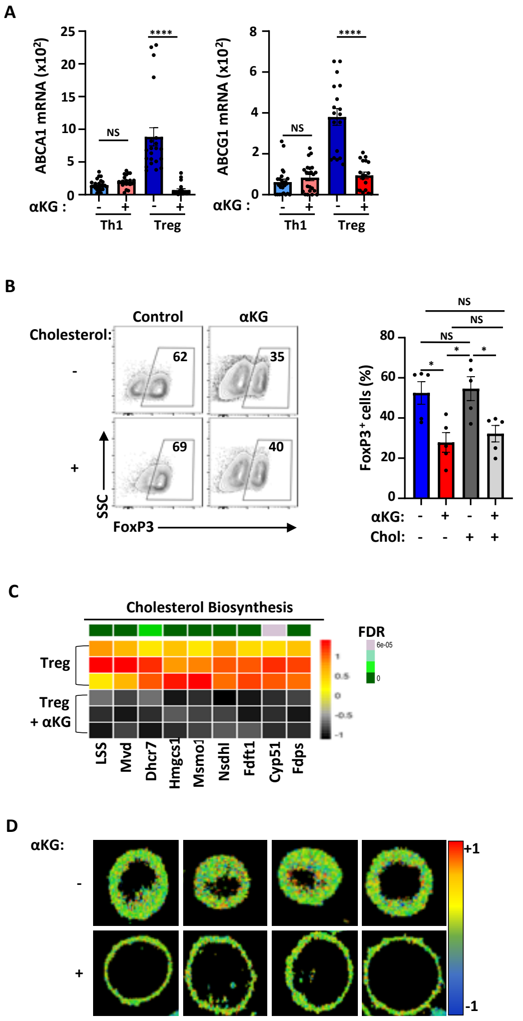


**(A)** *Abca1* and *Abcg1* transcripts were assessed by qRT-PCR and normalized to HPRT. Data are presented as means ± SEM of technical triplicates from 8 independent experiments. **(B)** Naïve CD4 T cells were activated in Treg polarizing conditions in the absence or presence of αKG and/or water-soluble cholesterol (50mM). FoxP3 expression was evaluated on day 4 by intracellular staining and representative dot plots are presented (left) as well as a quantification of the means ± SEM of 5 independent experiments (right). **(C)** A heatmap of RNASeq data showing differential expression of cholesterol biosynthesis genes in T cells activated in Treg-polarizing conditions in the absence or presence of αKG was created based on log2 transformed counts where each row represents an independent sample. **(D)** Representative generalized polarization (GP) images evaluating packing of membranes as a function of the red shift in loosely packed membranes. Statistical difference was determined by a one-way ANOVA and Tukey test for multiple comparisons (panel B) and an unpaired 2-tailed t-test (panel C; *, $p<0.05$; ****, $p<0.0001$; NS, not significant).

**Supplemental Figure 4: Decreased expression of fatty acid desaturases and elongation genes following Treg polarization in the presence of αKG.**

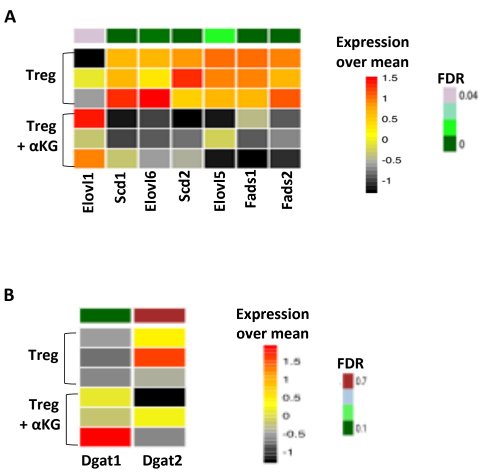


**(A)** A heatmap of RNASeq data for highly expressed fatty acid desaturases (*Scd1-3, Fads1-2*) and fatty acid chain elongation genes (*Elovl1, Elovl5, Elovl6*) in T cells activated in Treg-polarizing conditions in the absence or presence of αKG. Each row represents an independent sample. **(B)** A heatmap of RNASeq data for *Dgat1* and *Dgat2* as presented in panel A. Each row represents a single sample.

**Supplemental Figure 5: Utilization of glucose and glutamine carbons for the generation of TCA cycle intermediates is decreased in the presence of αKG.**

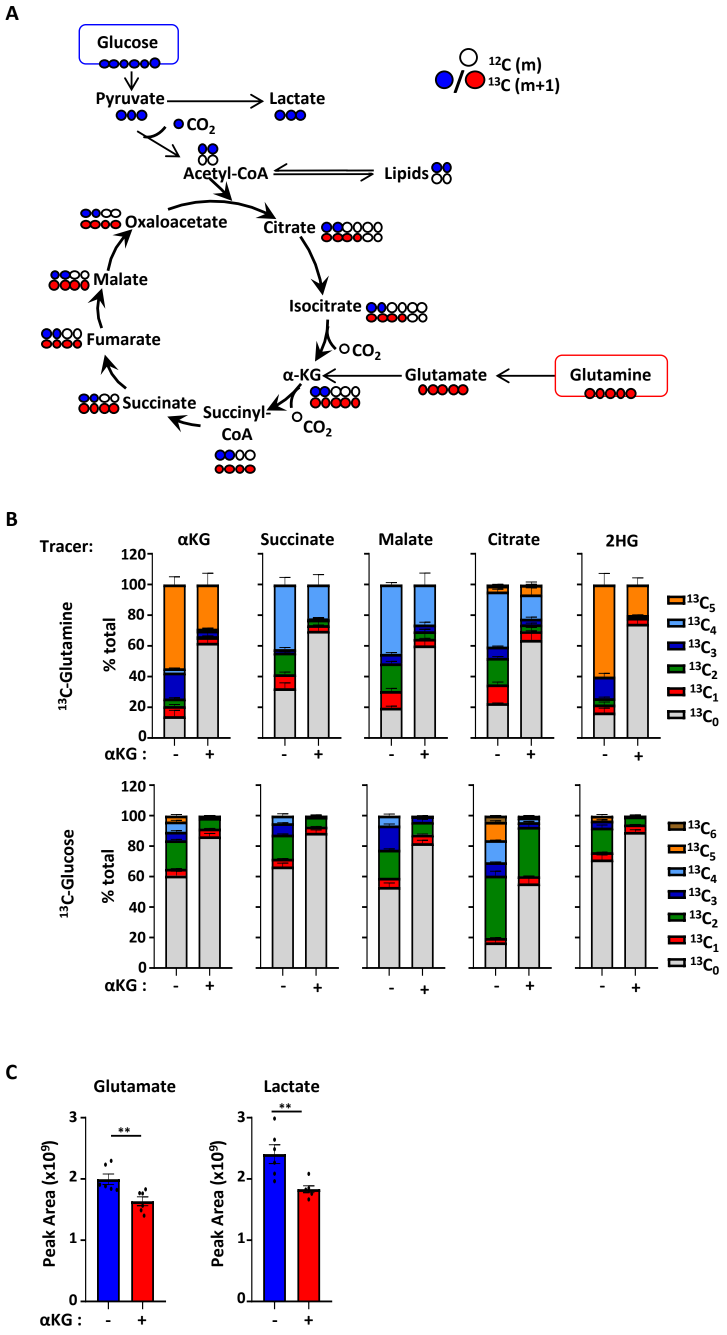


**(A)** Schematic representation of $^{13}C$ labelling products in the TCA cycle intermediates from [$^{13}C$]glucose and [$^{13}C$]glutamine. **(B)** The mean fractional abundance of the different isotopologues from [$^{13}C$]glutamine and [$^{13}C$]glucose into TCA cycle intermediates are shown as a percentage of the total. **(C)** The levels of glutamate and lactate were assessed in the indicated conditions by HPLC-MS and mean peak areas ± SEM are presented (n=6, 2 independent experiments). Statistical differences were determined by an unpaired 2-tail t-test (**, $p<0.01$; NS, not significant).

**Supplemental Figure 6: Exogenous succinate does not restore Treg polarization in the presence of αKG.**

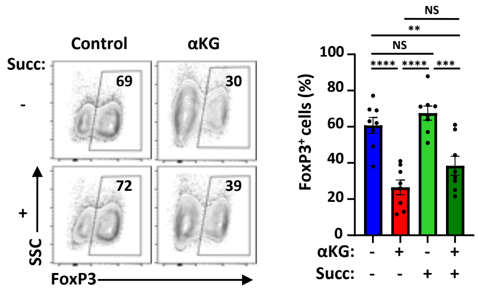


Naïve CD4 T cells were stimulated in Treg polarization conditions in the absence or presence of αKG and succinate as indicated (3.5 mM). FoxP3 expression was evaluated at day 4 by intracellular staining and representative plots are shown (left). Quantification of FoxP3 levels are presented as means ± SEM from 7 independent experiments and significance was evaluated by a one-way ANOVA and Tukey multiple comparison test (**, $p<0.01$; ***, $p<0.001$; ****, $p<0.0001$; NS, not significant).

**Supplemental Figure 7: Human CD4$^+$ T cells exhibit attenuated Treg polarization in the presence of αKG.**

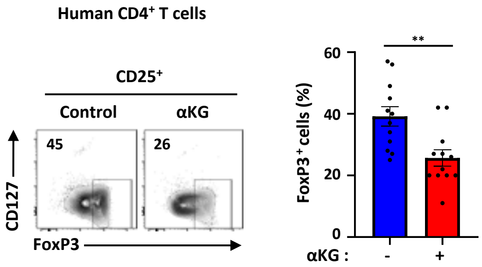


Naïve human CD4$^+$ T cells were activated in Treg-polarizing conditions and at day 4 of stimulation, Treg polarization was evaluated as a function of intracellular FoxP3 staining in gated CD25$^+$CD127$^{low}$ cells (left). Quantification of the percentages of FoxP3$^+$ cells is shown for 12 healthy donors (12 independent experiments) and significance was determined by an unpaired t-test (**, $p=0.003$).
