## Supplemental Tables for "αKG inhibits Regulatory T cell differentiation by coupling lipidome remodelling to mitochondrial metabolism"

### Supplemental Table 3: Murine Antibodies

| Antigen | Fluorochrome | Company | Clone | Catalog # |
|---|---|---|---|---|
| CD25 | AAF | EBIO | PC61.5 | 47-0251.82 |
| CD25 | PECY7 | BD | PC61 | 552880 |
| CD4 | BV711 | BD | RM4-5 | 563726 |
| CD4 | PE | BD | RM4-5 | 553049 |
| CD4 | PERCPCY5.5 | BD | RM4-5 | 550954 |
| CD4 | V450 | BD | RM4-5 | 560468 |
| CD4 | V650 | BD | RM4-5 | 563747 |
| CD44 | APCEF780 | EBIO | IM7 | 47-0441-82 |
| CD44 | PECY7 | EBIO | IM7 | 25-0441-82 |
| CD44 | V450 | BD | IM7 | 560451 |
| CD44 | V500 | BD | IM7 | 560780 |
| CD45 | APCEF780 | EBIO | 30-F11 | 47-0451-80 |
| CD45.2 | BV711 | BD | 104 | 563685 |
| CD45.2 | FITC | BD | 104 | 553772 |
| CD45.2 | PE | BD | 104 | 560695 |
| CD62L | APC | BD | MEL-14 | 553152 |
| CD62L | PE | BD | MEL-14 | 553151 |
| CD8α | FITC | BD | 53-6.7 | 553031 |
| CD8α | APC | BD | 53-6.7 | 553035 |
| CD8α | APC-R700 | BD | 53-6.7 | 564983 |
| CD8α | APCEF780 | EBIO | 53-6.7 | 47-0081-82 |
| CD8α | PERCPCY5.5 | BD | 53-6.7 | 551162 |
| FoxP3 | PECY7 | EBIO | FJK-16S | 25-5773-82 |

| | | | | |
|---|---|---|---|---|
| IFNγ | APC | EBIO | XMG1.2 | 17-7311-82 |
| T-bet | PE | EBIO | EBIO4B10 | 12-5825-82 |
| T-bet | PERCPCY5.5 | EBIO | EBIO4B10 | 45-5825-82 |
| T-bet | V650 | BD | O4-46 | 564142 |
| TCRβ | AF594 | BIOLEGEND | H57-597 | 109238 |

**Supplemental Table 4: Human Antibodies**

| Antigen | Fluorochrome | Company | Clone | Catalog # |
|---|---|---|---|---|
| CD4 | PercPCy5.5 | EBIO | OKT4 | 45-0048-42 |
| CD25 | FITC | Beckman Coulter | B1,49,9 | IM0478U |
| CD45RA | BV711 | BD | HI100 | 563733 |
| CD45RO | BV650 | BD | UCHL1 | 563750 |
| CD62L | PE | Beckman Coulter | DREG56 | IM2214U |
| CD62L | PE | BD | DREG56 | 555544 |
| CD127 (IL7Rα) | PECY7 | Beckman Coulter | R34.34 | A64618 |
| CD127 (IL7Rα) | BV786 | BD | HIL-7R-M21 | 563324 |
| FoxP3 | APC | EBIO | PCH101 | 17-4776-41 |

**Supplemental Table 5: Primer Sequences**

| Gene | Forward 5'-3' | Reverse 5'-3' |
|---|---|---|
| *foxp3* | GGCCCTTCTCCAGGACAGA | GCTGATCATGGCTGGGTTGT |
| *ifn*$\gamma$ | TGGCTCTGCAGGATTTTCATG | TCAAGTGGCATAGATGTGGAAGAA |
| *tbx21* | CAACAACCCCTTTGCCAAAG | TCCCCCAAGCAGTTGACAGT |
| *csf2* | TTTACTTTTCCTGGGCATTG | TAGCTGGCTGTCATGTTCAA |
| *tnf* | CATCTTCTCAAAATTCGAGTGACAA | GGGAGTAGACAAGGTACAACCC |
| *gzmb* | TGCTGACCTTGTCTCTGGCC | TAGTCTGGGTGGGGAATGCA |
| *abca1* | AAAACCGCAGACATCCTTCAG | CATACCGAAACTCGTTCACCC |
| *abcg1* | CTTTCCTACTCTGTACCCGAGG | CGGGGCATTCCATTGATAAGG |
| *hprt* | CTGGTGAAAAGGACCTCTCG | TGAAGTACTCATTATAGTCAAGGGCA |
